# Inhibiting RhoA Activation via GDP-state Stabilization to Relieve Heart Failure

**DOI:** 10.1101/2025.06.03.657556

**Authors:** Mengzhu Xue, Yingquan Liang, Zhen Yuan, Xiangning Liu, Longfeng Chang, Yongzhi Wang, Peijia Xu, Tingting Zhang, Hewei Jiang, Zijie Zhao, Jingqiu Liu, Shanshan Ruan, Tianyu Ye, Xuelian Pang, Wenyi Mei, Jiawen Wang, Xiaoqian Sun, Huijuan Wang, Jian Cui, Yao Zu, Xudong Lin, Zhenjiang Zhao, Rui Wang, Hong Huang, Cheng Luo, Shengce Tao, Jing Wang, Yajun Duan, Lili Zhu, Huifang Tang, Jian Zhang, Yong Wang, Chun Li, Honglin Li

## Abstract

High rates of heart failure (HF) morbidity and mortality have made targeting myocardial remodeling—particularly hypertrophy and fibrosis—a key therapeutic focus. RhoA, which regulates cytoskeletal reorganization and cell migration, plays a role in this process. However, RhoA has long been considered “undruggable”, due to its strong binding to its endogenous substrates, GDP/GTP, and the lack of well-defined pockets for drug targeting. Here, we discovered a cryptic pocket proximate to GDP within RhoA and identified a natural product, AH001, binds here and interacts with GDP, stabilizing RhoA’s interaction with its endogenous inhibitor, RhoGDIα. AH001 reduced the downstream MRTFA nuclear translocation and downregulated fibrosis/hypertrophy proteins. Consequently, AH001 mitigated myocardial remodeling in multiple HF animal models, and in the 3D myocardial tissue model. Our findings highlight the therapeutic potential of inhibiting RhoA activation in myocardial remodeling, ultimately targeting HF, and offer a promising avenue for developing reversible inhibitors against undruggable GTPases.

## INTRODUCTION

Heart failure (HF) remains the leading cause of global mortality, with rising incidence and persistently poor outcomes despite therapeutic advances.^1–3^ As a multi-faceted progressive syndrome, HF is driven by interconnected pathological processes, and it continues to impose a staggering global burden on more than 64 million affected individuals worldwide.^3, 4^ Myocardial remodeling, characterized by myocardial hypertrophy and fibrosis, is the most common and significant pathological process driving structural and functional cardiac deterioration in HF.^1, 4^ Despite advances in pharmacological therapies such as sodium-glucose cotransporter 2 inhibitors (SGLT2Is) and angiotensin receptor neprilysin inhibitors (ARNIs), the 5-year survival rate remains dismal at ∼50%, underscoring unmet clinical needs.^5–7^ The therapeutic gap stems from an incomplete understanding of the complexity of cellular and molecular mechanism in myocardial remodeling, such as structural changes and functional dysregulation of cells and proteins that occur at multiple levels.^8, 9^ Nevertheless, myocardial remodeling, characterized by pathologic hypertrophy and extracellular matrix (ECM) fibrosis, has been a cardinal independent predictor of HF progression based on emerging evidence.^10, 11^ How to reverse or prevent myocardial remodeling has become one of the important aspects in breaking through the bottleneck of HF treatments.

Ras homolog gene family member A (RhoA), is a small GTPase mediating physiological processes such as cytoskeletal reorganization and cell migration, and thereby involved in the pathogenesis of ischemia/reperfusion injury, hypertension, and heart failure.^12–14^ Regulation of RhoA occurs as a molecular switch between the RhoA-GDP (inactive) and RhoA-GTP (active) states.^15^ Once in the active RhoA-GTP state, RhoA signals allow the nuclear translocation of myocardin-related transcription factors (MRTFs) and stimulate serum response factor (SRF)-dependent transcription of fibrosis and hypertrophy-related genes.^16^ Therefore, inhibiting the activity of RhoA should be able to block the myocardial remodeling process. However, GTPases such as RhoA have a strong binding affinity for their endogenous substrates of GTP or GDP, which makes it difficult for drugs to displace these substrates.^17^ Additionally, RhoA rapidly cycles between the inactive GDP-bound and active GTP-bound states, which makes it difficult for drugs to stably bind to their active sites.^18^ Therefore, RhoA has long been considered as an “undruggable” target.

Here, we discovered a novel drug screening strategy targeting RhoA via a cryptic pocket, and identified a natural product, AH001. We performed surface plasmon resonance (SPR) to measure the binding affinity of AH001 for RhoA and determined the crystal structure of the RhoA-GDP-AH001 complex to elucidate its binding mode. We further elucidated the functional mechanisms of AH001 in fibroblasts, cardiomyocytes, and the 3D myocardial tissue model, and evaluated its efficacy in relieving myocardial remodeling in multiple HF models. Additionally, we performed the multi-omics data analysis demonstrating the association of RhoA activation and HF patients of diverse etiologies, and evaluated the level of RhoA activation by multiplex immunohistochemistry staining on sections of heart specimens from HF patients. Our work provides opportunities for the development of reversible inhibitors for RhoA, and establishes targeting RhoA as a previously unrecognized therapeutic strategy for myocardial remodeling in HF.

## RESULTS

### Activation of RhoA is associated with HF of diverse etiologies

Because *RHOA* showed gene-level associations with multiple cardiovascular diseases including coronary artery disease (CAD) and HF (Figures S1A and S1B), we investigated whether *RHOA* is associated with HF of diverse etiologies. We analyzed the cellular indexing of transcriptomes and epitomes by sequencing (CITE-seq) dataset and the single-nucleus RNA-sequencing (snRNA-seq) dataset on human left ventricle tissues (Figure 1A), including six healthy donors and 16 HF patients with acute myocardial infarction (AMI, n = 4), ischemic cardiomyopathies (ICM, n = 6) and non-ischemic cardiomyopathies (NICM, n = 6). These patients were diagnosed as HF with the average ejection fraction (EF) of 21.6 ± 10.02.^19^ Fibroblasts (40,370 cells) were separated from the CITE-seq dataset (Figure 1B) for further analysis. Differential gene expression (DGE) analysis exhibited that *RHOA* expression is higher in fibroblasts from AMI and NICM (Figure 1C). Specifically, the expression levels of *RHOA* and *MRTFA* were found to be positively correlated with the expression levels of the markers (Figure 1D), which were characteristics of disease-associated fibroblasts in HF.^19^ Cardiomyocytes (11,591 nuclei) were separated from the snRNA-seq dataset for further analysis (Figure 1E), no cardiomyocytes were obtained from the CITE-seq dataset. There is no significant difference in the overall expression levels of *RHOA* and *MRTFA* between the patients and healthy donors. Notably, the proportion of cardiomyocytes with moderate *RHOA* expression (UMI count >1) increased from 5% in healthy donors to 10-15% in patients with AMI, ICM, and NICM (Figure 1F). A similar trend was observed for *MRTFA* (10% in healthy donors increased to about 20% in patients with AMI, ICM, and NICM). Given the known roles of RHOA and MRTFA in fibrosis-related pathways,^13^ these *RHOA*+ or *MRTFA*+ subpopulations may represent stress-adapted cardiomyocytes that potentially respond to myocardial injury and contribute to disease progression.

**Figure 1.**
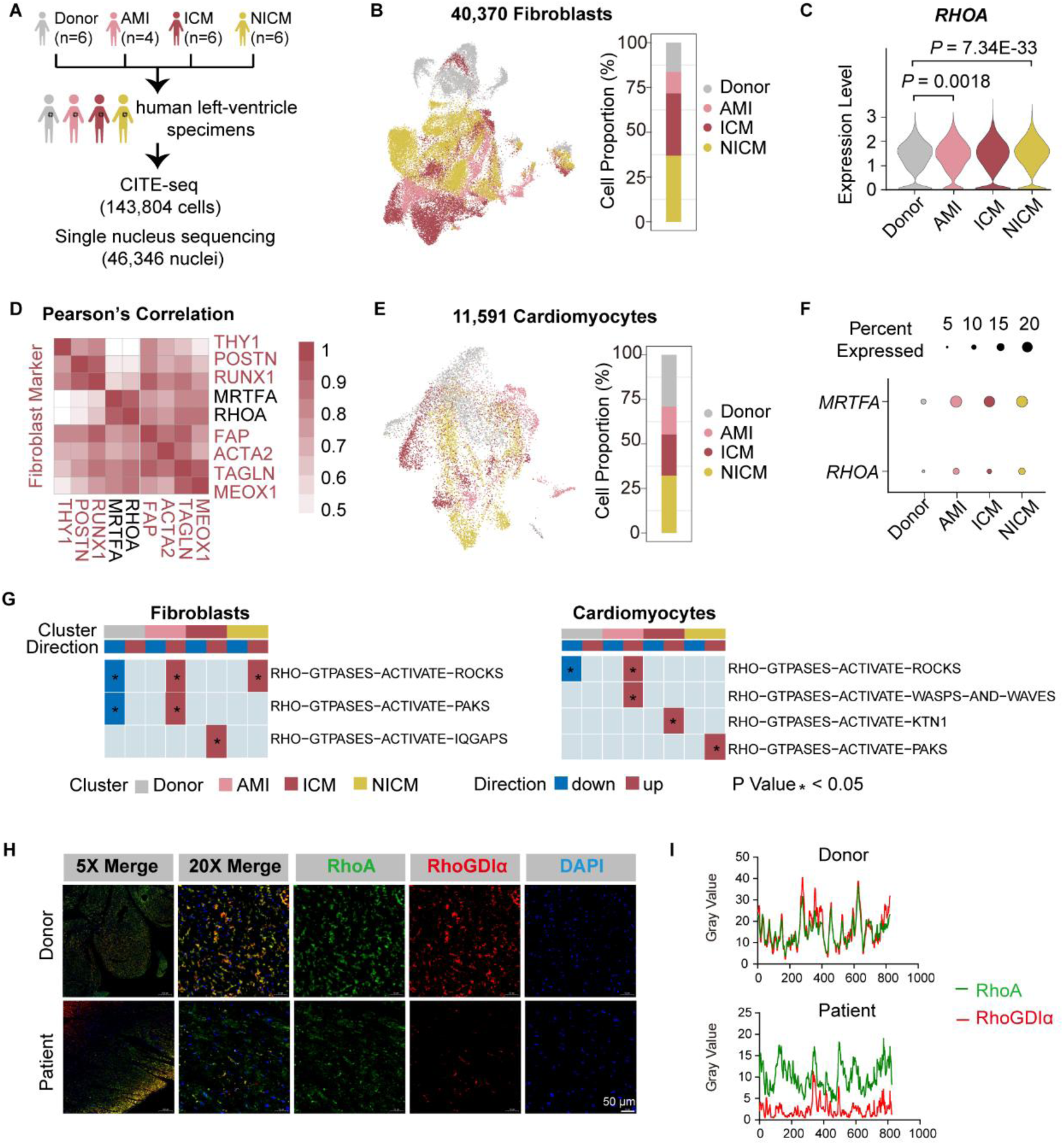
Activation of RhoA is associated with heart failure of diverse etiologies. (A) 22 specimens on human left ventricle tissues for the CITE-seq and snRNA-seq, including six healthy donors, four patients with AMI, six patients with ICM, and six patients with NICM.^19^ Ejection fraction (EF) is 21.6 ± 10.02 for patients. (B) 40,370 fibroblasts were separated from the CITE-seq, with cell proportions shown in donors and patients of AMI, ICM, and NICM. (C) The expression level of *RHOA* in fibroblasts from (B) as shown in donors and patients of AMI, ICM, and NICM. Adjusted p values were calculated by the DGE analysis. (D) Pearson’s correlation analysis of expression levels among selected genes in fibroblasts from donors and patients of AMI, ICM, and NICM. Markers that characterize disease-associated fibroblasts in heart failure were collected from the previous research ^19^ and labeled in red. (E) 11,591 cardiomyocytes were separated from the snRNA-seq, with cell proportions shown in donors and patients of AMI, ICM, and NICM. (F) Dotplot shows the percentage of cardiomyocytes expressing moderate *RHOA* (UMI count >1) or *MRTFA* (UMI count >1) in donors and patients of AMI, ICM, and NICM. (G) RHO GTPASE signaling enrichment analysis by irGSEA on fibroblasts and cardiomyocytes in donors and patients of AMI, ICM, and NICM. (H and I) The representative multiplex immunohistochemistry staining against RhoA and RhoGDIα on paraffin-embedded sections of interventricular septum (IVS) tissues from HF patients and healthy controls, with the quantification of colocalization of RhoA and RhoGDIα. Scale bar, 100 μm. See also Figure S1.

While our expression analysis revealed a disease-associated upregulation of *RHOA* in fibroblasts and a disease-associated expansion of *RHOA*+ or *MRTFA*+ cardiomyocytes, whether these phenotypes are driven by RhoA activation remains unclear. To evaluate the activation levels of RhoA, we extracted RHO-GTPASE signaling pathways from the REACTOME pathway database and performed the integrative robust gene set enrichment analysis (irGSEA) on the fibroblasts and cardiomyocytes. We found the activation levels of RHO-GTPASE signaling pathways were higher in the fibroblasts and cardiomyocytes from AMI, ICM, and NICM patients (Figure 1G). To confirm the results, we performed multiplex immunohistochemistry (mIHC) staining on paraffin-embedded sections of interventricular septum (IVS) specimens from HF patients and healthy donors (Figure S1C). The endogenous RhoA inhibitor RhoGDIα can bind to RhoA-GDP, preventing the release of GDP, and maintaining the complex integrity in the cytoplasm.^18^ We, therefore, quantified the colocalization of RhoA and the endogenous RhoA inhibitor RhoGDIα in the cytoplasm (Figure 1H). We observed fewer overlaps of the colocalization signals in the representative patient (Figure 1I). These results indicate that activation of RhoA is associated with HF of diverse etiologies.

### The drug screening strategy to stabilize RhoA in the inactivated GDP-bound state

RhoA lacks well-defined binding pockets for compounds due to the conformational flexibility resulting from the GTP/GDP binding, except for the compound binding pocket that covalently binds to cysteine 107.^20^ To systematically analyze the conformational flexibility of RhoA in active and inactive states, we performed extensive atomic-level simulations on structures of RhoA-GTP (PDB: 1A2B) and RhoA-GDP (PDB:1FTN) (Figures S2A and S2B). Compared to the GTP-bound state, the absence of the γ-phosphate in GDP increases the flexibility of the switch regions of RhoA-GDP, facilitating conformational changes and unveiling a cryptic pocket that opens dynamically during simulations (Figure S2C). The cryptic pocket is located near the GDP and extends to switch I (SW-I) and switch II (SW-II) regions in the RhoA-GDP structure (Figure 2A). We, therefore, propose a strategy to stabilize RhoA in the inactivated GDP-bound state by binding a compound at the pocket (Figure 2A).

**Figure 2.**
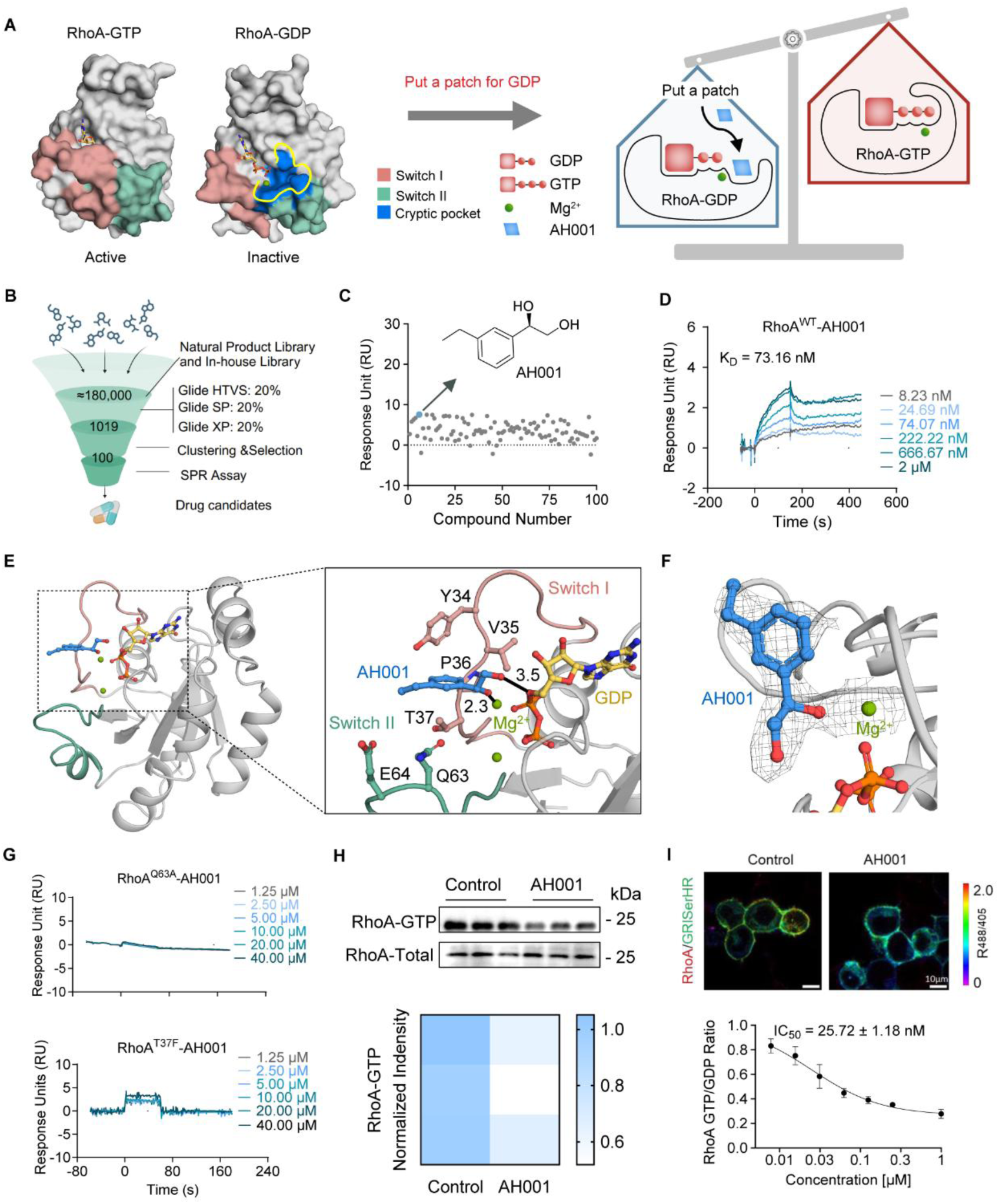
A new strategy to identify a natural product restricting RhoA in the GDP-bound state. (A) The scheme for the “put a patch” drug screening strategy targeting RhoA. The left part illustrated the identification of a cryptic pocket located near GDP from the structural difference analysis, shown with RhoA-GTP (PDB: 1A2B) and RhoA-GDP (PDB: 1FTN). The right part illustrated the drug screening strategy that put a compound at the cryptic pocket to act as a patch for GDP, stabilizing the GDP binding and inhibiting GTP competition. (B and C) Virtual screening of a natural compound library and in-house library using this “put a patch” strategy (B), followed by SPR assays to obtain AH001 as the potential drug candidate (C). (D) The dissociation constant (K_D_) of AH001 for RhoA was determined using SPR assays. (E) Co-crystal structure of AH001 (blue) and RhoA (green) with GDP (yellow) and Mg^2+^ (dark green) showed switch I and switch II, with the zoomed view of AH001 in the binding pocket. The black dashed lines represented the distance (in Å) between heavy atoms involved in a coordination bond or hydrogen bonds. (F) The 2Fo-Fc electron density map of compound AH001 with Mg^2+^. (G) The dissociation constant (K_D_) of AH001 for RhoA^Q63A^ and RhoA^V38A^ mutants was determined using SPR assays. (H) The expression level of RhoA-GTP was detected by the pull-down assay in HEK 293F cells treated with AH001 at 20 μM, with the normalized intensity shown in the heatmap. (I) RhoA GTP/GDP ratio was evaluated by a fluorescent probe in HEK 293F cells, with the treatment of AH001 at 1 μM. IC_50_ values were inferred by RhoA GTP/GDP ratios in HEK293F cells, with the treatment of AH001 (n = 3) at doses of 7.8 nM to 1 μM. Scale bar, 10 μm. Statistical analysis was performed using one-way analysis of variance (ANOVA) among multiple groups, which was followed by an LSD post hoc test between two groups. See also Figure S2 and Table S1.

### Identification of a natural product via the drug screening strategy

Based on the drug screening strategy, we performed a compound virtual screening assay from the TOPSCIENCE natural product library and an in-house library (∼180,000 compounds in total). The top 1,019 compounds were prioritized for further evaluation based on binding mode, structural diversity, and novelty. From these, 100 candidates were selected for surface plasmon resonance (SPR) assays to assess their binding affinity with RhoA (Figure 2B). AH001, a natural product with excellent binding mode and binding affinity with RhoA, was selected from the screening, showing an equilibrium dissociation constant (K_D_) of 73.16 nM to RhoA (Figures 2C and 2D). To verify the effectiveness of our strategy and characterize the binding mechanism of AH001 to RhoA-GDP, we obtained the co-crystal structure of RhoA-GDP/AH001 complex at a resolution of 3.1 Å (PDB: 9J0G; Table S1). As shown in the crystal structure, AH001 is located in the cavity formed by SW-I and SW-II, and the site is overlapped with the cryptic pocket identified by structural analysis (Figure 2E). The phenethyl group of AH001 bound in the hydrophobic cleft surrounded by the Y37, V35, P36, T37, and Q63 residues (Figure 2E). Specifically, AH001 formed a coordination bond with Mg^2+^ and a hydrogen bond with GDP (Figure 2F). The binding affinity of AH001 to RhoA was diminished in both the T37F and the Q63A mutants (Figure 2G). To further reveal structural changes induced by AH001, we aligned the ligand-free (RhoA-GDP) and ligand-bound RhoA (RhoA-GDP-AH001) structures. Significant conformational changes were observed in the switch regions, particularly in SW-II. The Cα atoms of D65, Y66, and D67 in RhoA-GDP-AH001 were shifted away from the positions of the corresponding residues in RhoA-GDP with the distances ranging from ∼ 2.0 to 4.6 Å (Figure S2D).

To further evaluate the inhibitory effects of AH001 on RhoA, we conducted a series of functional assays. Firstly, we observed reduced levels of RhoA-GTP in HEK 293F cells (Figure 2H). Secondly, we used a fluorescent probe to calculate the fluorescent intensity ratio of F488 nm/F405 nm to evaluate the cellular RhoA GTP/GDP exchange ratio (Figures S2E and S2F). A concentration-dependent decrease in the RhoA GTP/GDP exchange ratio was observed under treatments of AH001 at 7.8 nM to 1 μM, resulting in an IC_50_ value of 25.72 ± 1.18 nM (Figure 2I). Lastly, we investigated whether AH001 would affect the binding of RhoA to its endogenous inhibitor RhoGDIα. Through structural analysis, we found that in the presence of AH001, the shifted SW-II region displayed significant spatial complementarity with a “regulatory arm” located at the N terminus (residues 5-55) of RhoGDIα, which may facilitate the interaction between RhoA and RhoGDIα (Figure S2G). To investigate whether AH001 can facilitate the RhoA-RhoGDIα binding through the “regulatory arm” of RhoGDIα, we generated a truncated RhoGDIα protein with the deletion of the “regulatory arm” (Figure S2H). The interaction between RhoA and RhoGDIα was significantly decreased when the “regulatory arm” of RhoGDIα was truncated, as demonstrated by the proximity ligation assay (PLA) (Figure S2I). These results confirm that AH001 can interact with RhoA-GDP, stabilize the RhoA-RhoGDIα binding to constrain RhoA in the GDP-bound state.

### AH001 alleviates myocardial remodeling in multiple HF animal models

To evaluate the in vivo efficacy of AH001, we first characterized its pharmacokinetic profile and conducted systemic safety assessments, confirming the absence of acute toxicity across multiple dose levels in various animal species (Figures S3A and S3B). Then, we established multiple HF animal models to investigate whether AH001 can alleviate myocardial remodeling. To stimulate myocardial hypertrophy and fibrosis in HF, we first established ISO-induced cardiomyocyte hypertrophy models in zebrafish at 2 dpf, treated with AH001 or LCZ696 (an ARNI, as the positive control) (Figure 3A). AH001 administration improved the ISO-induced myocardial hypertrophy, the ejection fraction (EF), the fractional shortening (FS), the blood flow rate, ventricular area, and cardiomyocyte area in zebrafishes with the minimum dose of 15.16 μM (Figures 3B-3D), which is comparable to LCZ696 at the dose of 69.59 μM. To further evaluate the efficacy of AH001 on improving myocardial hypertrophy and fibrosis, we established Angiotensin II (Ang II)-induced models in 8-week-old mice (Figure S4A). After four weeks of oral gavage administration, mice treated with AH001 at 10 mg/kg showed improved cardiac functions comparable to the mice treated with LCZ696 at 66.67 mg/kg, assessed by echocardiography, Masson’s trichrome staining, and wheat germ agglutinin (WGA) staining (Figures S4B-S4F). Consistently, elevated EF and FS, reduced fibrosis areas, and decreased cardiomyocyte areas were observed in mice treated with AH001 or LCZ696. Both the systolic blood pressure (SBP) and the diastolic blood pressure (DBP) of Ang II-induced mice were restored to normal levels after AH001 or LCZ696 treatments (Figure S4G).

**Figure 3.**
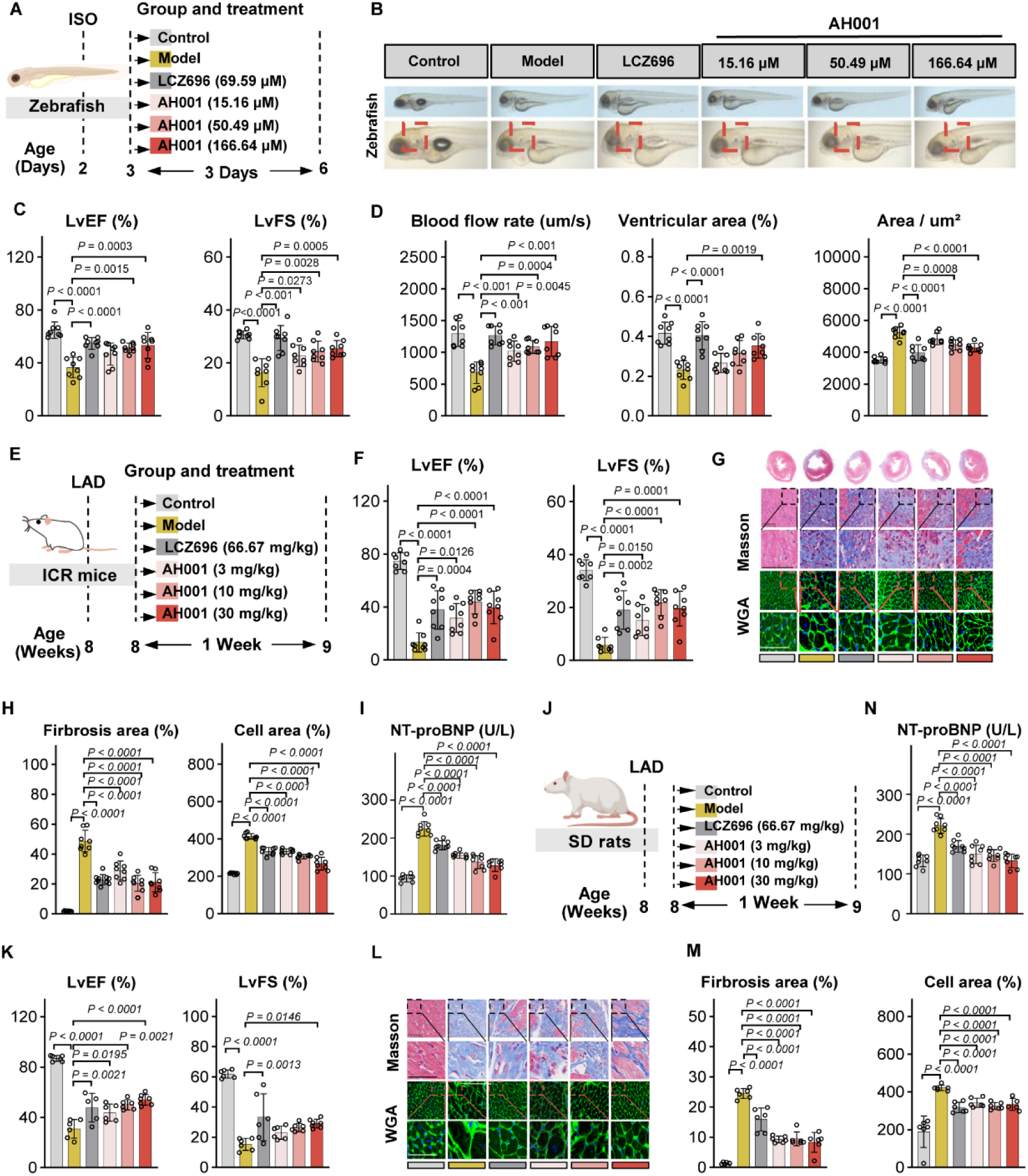
AH001 alleviated myocardial remodeling in multiple animal models. (A) The schema of ISO-induced myocardial hypertrophy zebrafish models (n = 10) construction, followed by treatments of LCZ696 (69.59 μM) and AH001 at doses of 15.16, 50.49, and 166.64 μM. (B) The representative images of hearts from the ISO-induced zebrafish models, followed by treatments of LCZ696 and AH001 described in (A). (C) EF and FS in ISO-induced zebrafish models by the Echocardiographic assessment. (D) Blood flow rate, ventricular area, and cardiomyocyte area in ISO-induced zebrafish models. (E) The schema of LAD mouse models (n = 8) construction, followed by treatments of LCZ696 (66.67 mg/kg) and AH001 at doses of 3, 10, and 30 mg/kg. (F) EF and FS in LAD mouse models by the Echocardiographic assessment. (G and H) Representative images of Masson’s trichrome staining and WGA staining of a cross-sectional heart specimen, shown in the left ventricle region (G). Scale bar, 1 mm. Cardiac fibrosis was assessed by the fibrosis area and the cross-sectional area of cardiomyocytes (H). (I) Determination of NT-proBNP in serum from LAD mouse models. (J) The schema of LAD rat models (n = 6) construction, followed by treatments of LCZ696 (66.67 mg/kg) and AH001 at doses of 3, 10, and 30 mg/kg. (K) EF and FS in LAD rat models by the Echocardiographic assessment. (L and M) Representative images of Masson’s trichrome staining and WGA staining of a cross-sectional heart specimen, shown in the left ventricle region (L). Scale bar, 1 mm. Cardiac fibrosis was assessed by the fibrosis area and the cross-sectional area of cardiomyocytes (M). (N) Determination of NT-proBNP in serum from LAD rat models. Data are mean ± SEM. Statistical analysis was performed using one-way analysis of variance (ANOVA) among multiple groups, which was followed by an LSD post hoc test between two groups. See also Figure S3, Figure S4, and Figure S5.

To stimulate myocardial infarction, we established LAD models in 8-week-old mice (Figure 3E) and assessed cardiac functions by echocardiography, Masson’s trichrome staining, and WGA staining. Masson’s trichrome staining showed a significant increase in blue-stained collagen fiber area in the heart tissue from LAD mouse models, suggesting enhanced collagen deposition and fibrosis formation. WGA signal at the cardiomyocyte membrane appeared widened in LAD mouse models, suggesting an increase in cell surface area. After one week of oral gavage administration, the mice treated with AH001 showed improved cardiac functions at the minimum dose of 3 mg/kg, as shown in Figures 3F-3H, which is comparable to LCZ696 at a 66.67 mg/kg dose. To assess the extent of myocardial injury, we determined levels of NT-proBNP, lactate dehydrogenase (LDH), creatine kinase (CK), and Ang II in the serum of LAD mouse models, all of which showed significant reductions after AH001 treatments (Figures 3I and S4H). We established LAD models in 8-week-old rats for further assessments (Figure 3J). Consistently, AH001 showed improved cardiac functions (Figures 3K-3M). Specifically, AH001 significantly reduced the levels of NT-proBNP, LDH, CK, and BNP in the serum of LAD rat models (Figures 3N and S4I). Lastly, we established a doxorubicin (Dox)-induced cardiotoxicity model in 8-week-old mice to simulate HF (Figure S5A). Although AH001 did not improve the body weight loss in Dox-induced models, AH001 significantly improved cardiac functions and the extent of myocardial injury (Figures S5C-S5H). Hematoxylin and Eosin (HE) staining and Sirius red staining results revealed that AH001 improved cardiac vacuole index and fibrosis in the heart (Figures S5I-S5L). These results confirm that AH001 alleviates myocardial remodeling in multiple HF animal models.

### AH001 simultaneously targets both fibroblasts and cardiomyocytes via MRTFA-associated pro-fibrotic pathways

To further understand how AH001 alleviates myocardial remodeling, we performed single-cell RNA-sequencing (scRNA-seq) in heart specimens obtained from LAD mouse models that were treated with oral gavage administration of AH001 at 10 mg/kg for 7 days (Figure 4A). We obtained 105,123 cells in total with 27,838 fibroblasts separated (Figure 4B), and observed the upregulated expression of *Rhoa* in LAD mouse models, which was reversed by the AH001 treatment (Figure 4C). Consistent with the positive expression correlation between *RHOA* and disease-associated fibroblast markers shown in Figure 1D, we observed a notable upregulation of *Postn*, *Acta2*, and *Runx1* in LAD mouse models, which was reversed by the AH001 treatment (Figure 4C). We also separated 56,081 cardiomyocytes (Figure 4D), and observed the downregulated expression of *Rhoa* and *Nppa* (Precursor of ANP, a marker for HF) in LAD mouse models treated by AH001 (Figure 4E). Although no significant intergroup difference in overall *Mrtfa* expression was observed in cardiomyocytes, it is noteworthy that approximately 20% of cardiomyocytes consistently exhibited moderate *Mrtfa* expression levels (UMI count >1) in our mouse models (Figure 4F).

**Figure 4.**
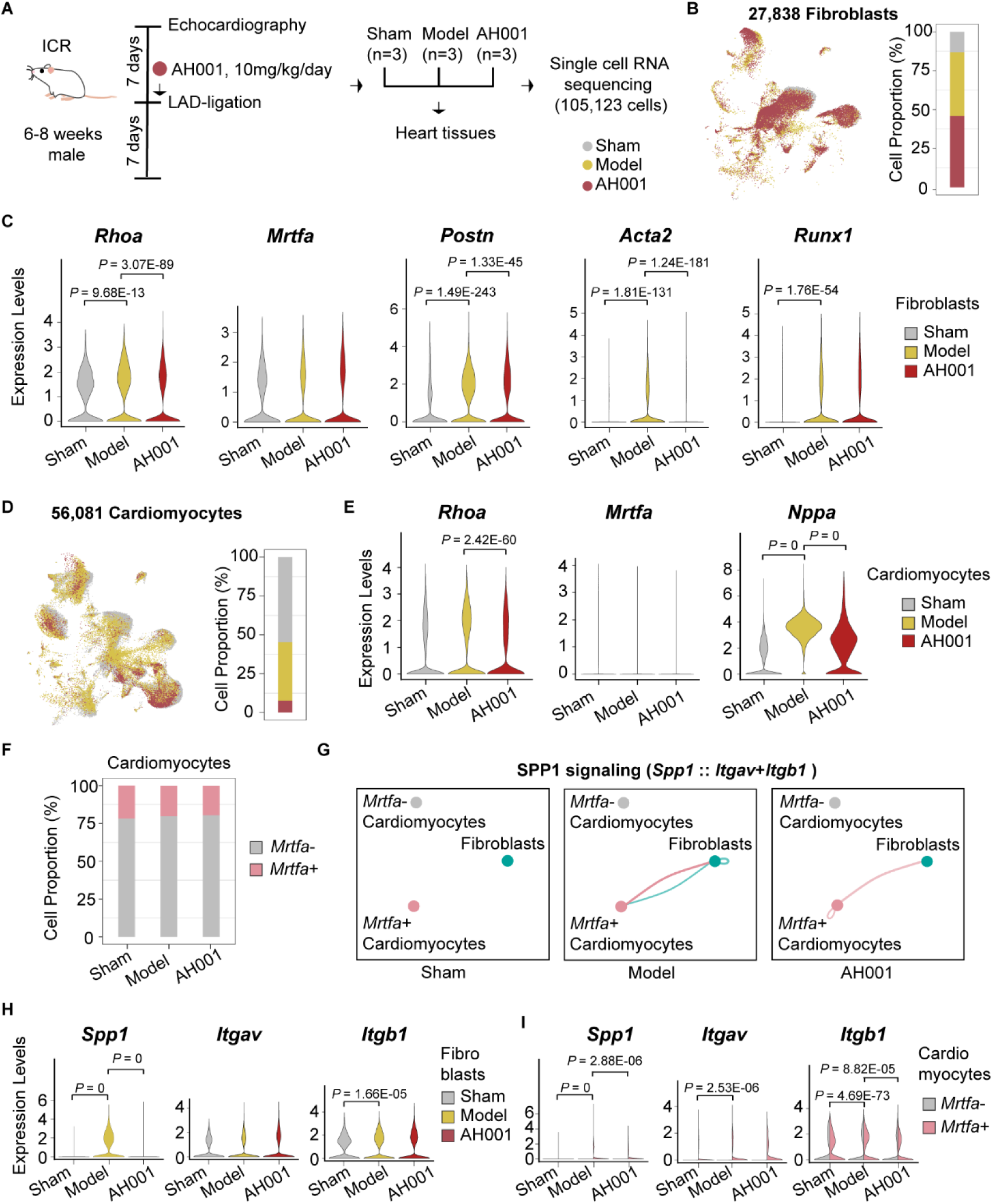
The scRNA-seq analysis revealed that AH001 impacted fibroblasts and cardiomyocytes. (A) The schema of heart specimens’ collection from LAD mouse models (n = 3), followed by the treatment of AH001 at 10 mg/kg for seven days. A total of 105,123 cells were obtained from the scRNA-seq analysis. (B) 27,838 fibroblasts were separated from the scRNA-seq, with cell proportions shown in Sham, Model, and AH001 groups. (C) Expression levels of *Rhoa*, *Mrtfa,* and disease-associated fibroblast markers from Figure 1D of *Postn*, *Acta2*, and *Runx1* in fibroblasts from (B) were shown in Sham, Model, and AH001 groups. Adjusted p values were calculated by the DGE analysis. (D) 56,081 cardiomyocytes were separated from the scRNA-seq, with cell proportions shown in Sham, Model, and AH001 groups. (E) Expression levels of *Rhoa*, *Nppa* (a HF marker), and *Mrtfa* in cardiomyocytes from (D) (D) were shown in Sham, Model, and AH001 groups. Adjusted p values were calculated by the DGE analysis. (F) Cell proportions of *Mrtfa+* and *Mrtfa-* cardiomyocytes were shown in Sham, Model, and AH001 groups. (G) SPP1 signaling between *Mrtfa+*, *Mrtfa-* cardiomyocytes, and fibroblasts were shown in Sham, Model, and AH001 groups, with the biggest contribution from the ligand-receptor pair of *Spp1* to *Itgav* and *Itgb1*. (H and I) Expression levels of *Spp1*, *Itgav*, and *Itgb1* in fibroblasts (H), *Mrtfa+*, and *Mrtfa-*cardiomyocytes (I) were shown in Sham, Model, and AH001 groups. Adjusted p values were calculated by the DGE analysis.

Considering the disease-associated expansion of *MRTFA*+ cardiomyocytes in HF patients (Figure 1F) and the established role of MRTFA in fibrosis-related pathways,^13, 21^ we hypothesized that these *Mrtfa+* cardiomyocytes may contribute to maladaptive remodeling through paracrine signaling in response to myocardial injury. To verify this hypothesis, we divided the cardiomyocytes into *Mrtfa*+ and *Mrtfa*-subpopulations according to the expression level of *Mrtfa*, and investigated whether there were differences in the cardiomyocyte-fibroblast interactions between the two subpopulations. We found that *Mrtfa+* cardiomyocytes could interact with fibroblasts mutually through the SPP1 signaling (mediating pathological hypertrophy and progressive fibrosis) ^22, 23^ in LAD mouse models, with the biggest contribution from the ligand-receptor pairs of *Spp1* to *Itgav* and *Itgb1* (Figure 4G). Notably, *Spp1* (Osteopontin, OPN) expression was significantly upregulated in fibroblasts from LAD mouse models, which was reversed by AH001 treatment (Figure 4H). This expression pattern was specifically recapitulated in *Mrtfa+* cardiomyocytes, whereas *Mrtfa*-cardiomyocytes showed negligible *Spp1* expression (Figure 4I). Interestingly, while *Spp1* expression was regulated in both fibroblasts and *Mrtfa+* cardiomyocytes, its receptor *Itgb1* exhibited distinct cell type-specific regulation. *Itgb1* expression was significantly upregulated in LAD mouse models and downregulated by AH001 treatment exclusively in *Mrtfa+* cardiomyocytes (Figure 4I). These results suggest MRTFA as a marker of a cardiomyocyte subpopulation with unique fibrotic cell-cell interaction potential, and indicate that SPP1 signaling serves as a key mediator of myocardial hypertrophy and fibroblast activation.

### AH001 inhibits the downstream signaling of RhoA activation in vitro

The significant manifestation of RhoA activation is the nuclear translocation of MRTFA, while the indication of RhoA inactivation is the formation of RhoA-RhoGDIα complex (Figure 5A). To investigate whether AH001 suppressed fibroblast activation via the MRTFA nuclear translocation mechanism, we established a cardiac fibroblast activation model (Figure 5B). We isolated cardiac fibroblasts from neonatal rats for in vitro culture, and treated them with TGF-β1 (to induce fibroblast activation) and AH001 at a dose series of 10, 20, and 40 μM (Figure 5B). AH001 effectively inhibited the migration of fibroblasts, as demonstrated by wound healing and Transwell assays (Figures 5C and 5D; S6A and S6B). AH001 also reduced the level of RhoA-GTP, the F-actin formation, and the nuclear translocation of MRTFA (Figures 5E-5G). Consistently, we observed decreased levels of fibrosis-related proteins in fibroblasts after the AH001 treatment, including Fibronectin 1 (FN1), Collagen type III (COL3), Serum Response Factor (SRF), Vimentin (VIM), and the phosphorylation of Cofilin (CFN) (Figure S6C). Further, RhoGDIα knockdown attenuated the AH001-induced reductions in the F-actin formation and the nuclear translocation of MRTFA (Figures 5H and 5I). These results suggest that AH001 acts on RhoA-RhoGDIα axis to inhibit the downstream signaling of RhoA activation in fibroblasts.

**Figure 5.**
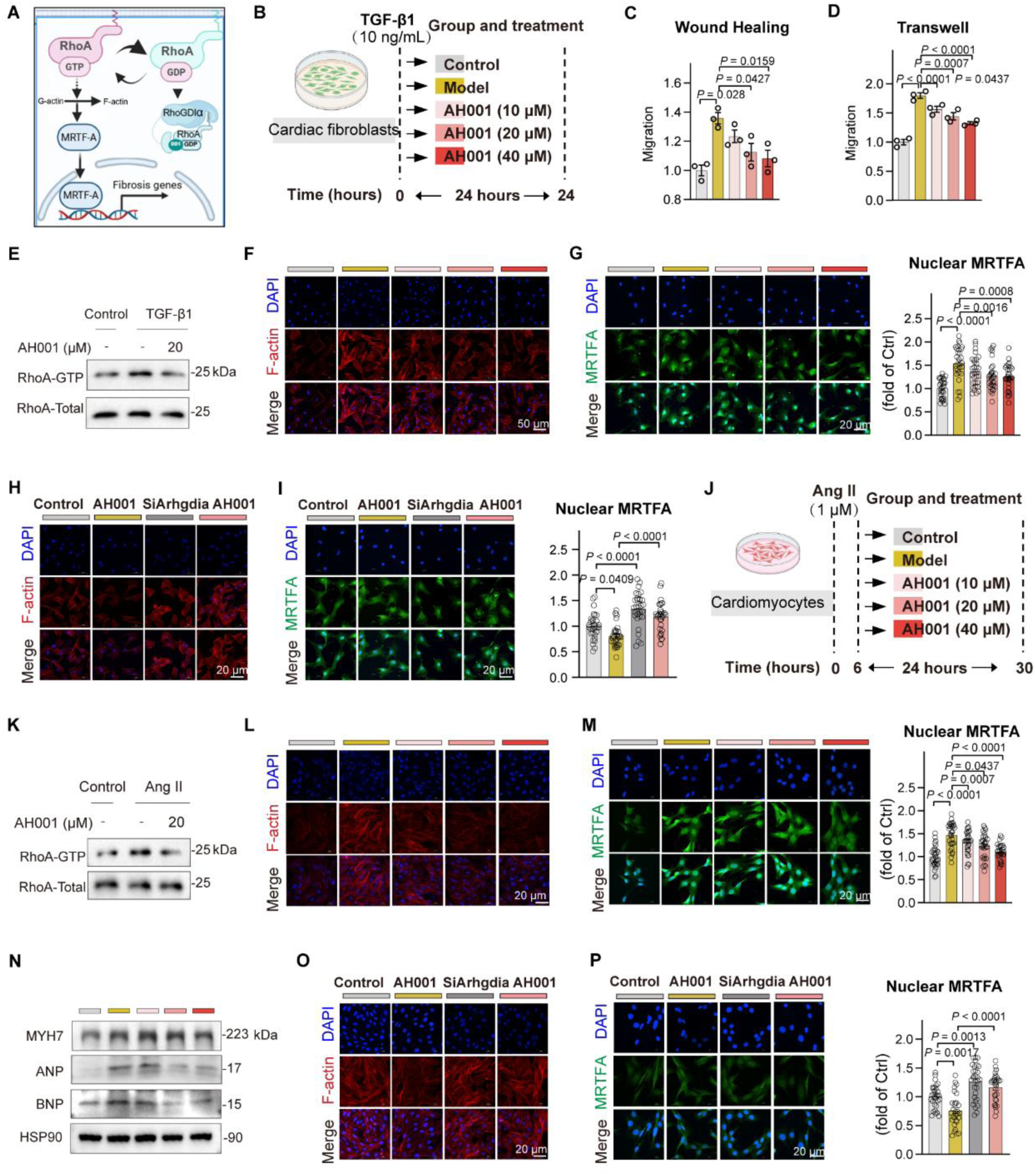
AH001 suppressed the nuclear translocation of MRTFA in fibroblasts and cardiomyocytes. (A) The schema of canonical downstream signaling of RhoA activation and inactivation. (B) The schema of fibroblast activation cellular model construction, followed by treatments of AH001 at doses of 10, 20, and 40 μM. (C and D) The impact of AH001 on the migration of fibroblasts (n = 3) as assessed by the wound healing assay and the Transwell assay. (E) The expression level of RhoA-GTP was detected by the pull-down assay in fibroblasts with the treatment of AH001 at 20 μM. (F and G) The representative immunofluorescence staining against F-actin (F) and MRTFA (G) in fibroblasts, with the quantification of nuclear MRTFA fluorescence signals. Scale bar, 50 μm (F), 20 μm (G). (H and I) The representative immunofluorescence staining against F-actin (H) and MRTFA (I) in fibroblasts followed by the knockdown of RhoGDIα and the treatment of AH001 at 20 μM, with the quantification of nuclear MRTFA fluorescence signals. Scale bar, 20 μm. (J) The schema of the myocardial hypertrophy cellular model construction, followed by treatments of AH001 at doses of 10, 20, and 40 μM. (K) The expression level of RhoA-GTP was detected by the pull-down assay in cardiomyocytes with the treatment of AH001 at 20 μM. (L and M) The representative immunofluorescence staining against F-actin (L) and MRTFA (M) in cardiomyocytes, with the quantification of nuclear MRTFA fluorescence signals. Scale bar, 20 μm. (N) Expression levels of proteins involved in myocardial hypertrophy determined by WB, including MYH7, ANP, BNP, and HSP90. (O and P) The representative immunofluorescence staining against F-actin (O) and MRTFA (P) in cardiomyocytes followed by the knockdown of RhoGDIα and the treatment of AH001 at 20 μM, with the quantification of nuclear MRTFA fluorescence signals. Scale bar, 20 μm. Data are mean ± SEM. Statistical analysis was performed using one-way analysis of variance (ANOVA) among multiple groups, which was followed by an LSD post hoc test between two groups. See also Figure S6.

Next, we investigated whether AH001 inhibited myocardial hypertrophy via this mechanism. We isolated cardiomyocytes from neonatal rats for in vitro culture, and treated with Ang II (to induce myocardial hypertrophy) and AH001 at a dose series of 10, 20, and 40 μM (Figure 5J). AH001 effectively reduced myocardial hypertrophy, as measured by the fluorescent area of ACTN2 staining (Figure S6D). Similarly, we observed a decrease in the level of RhoA-GTP, F-actin formation, nuclear translocation of MRTFA, and fibrosis-related proteins after the AH001 treatment (Figures 5K-5M and S6E). Additionally, decreased levels of proteins involved in myocardial hypertrophy were observed after the AH001 treatment, including MYH7, ANP, and BNP (Figure 5N). After the knockdown of RhoGDIα in cardiomyocytes, the effects on the F-actin formation, the nuclear translocation of MRTFA, and the fluorescent area of ACTN2 staining by the AH001 treatment were decreased (Figures 5O-5P and S6F). These results suggest that AH001 acts on RhoA-RhoGDIα axis to inhibit the downstream signaling of RhoA activation in cardiomyocytes.

Considering that AH001 downregulated the levels of secreted pro-fibrotic proteins such as FN1, COL3, and VIM in cardiomyocytes (Figure S6E), we investigated whether fibroblast activation could be affected by in vitro culture with the conditional medium from cardiomyocyte culture. We established an in vitro culture model of cardiomyocytes under anaerobic conditions (AC), followed by treatments of AH001 at a dose series of 10, 20, and 40 μM (Figure S6G). AH001 reduced the levels of secreted pro-fibrotic factors, such as SPP1, Connective Tissue Growth Factor (CTGF, a fibrosis marker), and TGF-β in the culture supernatant (Figure S6H). Using the conditional medium from cardiomyocyte culture, we observed increased migration of fibroblasts in the model group and decreased migration of fibroblasts in the AH001-treated groups by the Transwell assay (Figure S6I). These results demonstrate that cardiomyocyte-derived secreted factors promote fibroblast activation, and AH001 effectively suppress this paracrine-mediated pro-fibrotic response.

### AH001 improved fibrosis in the 3D myocardial tissue model

To further evaluate the dual regulation of AH001 to cardiomyocytes and fibroblasts, we prepared a 3D myocardial tissue model under anaerobic conditions (Figure 6A), thus providing visual and quantitative results for drug evaluation.^24^ Co-staining with α-actin (a cardiomyocyte marker) and vimentin (a fibroblast marker) antibodies enables visualization of cardiomyocyte-fibroblast intermingling in the 3D myocardial tissue model (Figure 6B). The decreased colocalization of RhoA and RhoGDIα was also observed in the 3D myocardial tissue model (Figures 6C and 6D), demonstrating the exceptional biomimetic fidelity of this model in recapitulating failing human myocardium. In the model group, cardiomyocytes exhibited significant pathological contraction, with their diameter reduced by more than half (Figure 6E). Concurrently, the fluorescent area of cardiomyocytes diminished to approximately 10% (Figure 6F). By contrast, the proliferation of fibroblasts in the model group increased by 170% via the cell viability assay (Figure 6G). It indicates that the reduction in the size of the cell spheroids is not caused by cardiomyocyte necrosis, but rather due to the proliferation of fibroblasts. AH001 effectively suppressed the proliferation of fibroblasts, thereby significantly alleviating the pathological contraction of cardiomyocytes in a dose-dependent manner. To further verify the activation of fibroblasts, we assessed the levels of fibrotic makers of α-SMA (alpha-smooth muscle actin) and Vimentin by immunofluorescence (Figure 6H). The fluorescent intensity of α-SMA and Vimentin increased by nearly 10-fold in the model group compared to the control group (Figure 6I). AH001 significantly reduced the expression levels of α-SMA and Vimentin, assessed by the fluorescent signals and the fluorescent intensity of these proteins. We observed that AH001 decreased the levels of pro-fibrotic factors such as COL1A1, COL3A1, SPP1, and Galectin3 in the culture supernatant in a dose-dependent manner (Figure 6J), suggesting AH001 inhibition of fibroblast activation.

**Figure 6.**
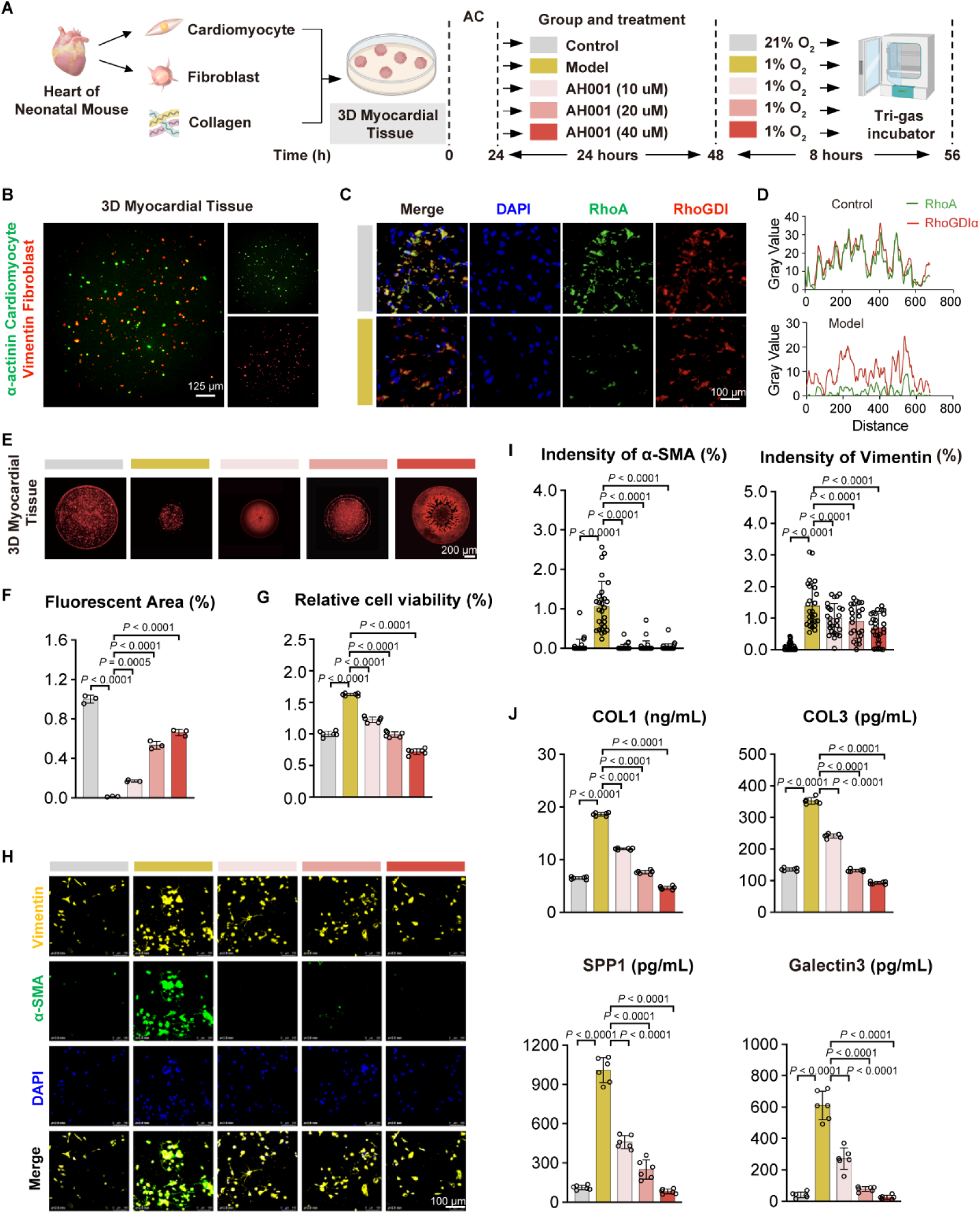
The impacts of AH001 on the 3D myocardial tissue model. (A) The schema of the 3D myocardial tissue model construction integrating cardiomyocytes and fibroblasts, followed by treatments of AH001 at doses of 10, 20, and 40 μM. (B) The representative live immunofluorescence staining against α-actinin (cardiomyocyte marker) and Vimentin (fibroblast marker) in the 3D myocardial tissue model. Scale bar, 125 μm. (C and D) The representative multiplex immunohistochemistry staining against RhoA and RhoGDIα on paraffin-embedded sections of the 3D myocardial tissue model (C), with the quantification of colocalization of RhoA and RhoGDIα (D). Scale bar, 100 μm. (E and F) The representative images (E) for the contractile phenotype of AC-induced fibrosis in the 3D myocardial tissue model, with the fluorescent areas detected using Image J (F). Scale bar, 200 μm. (G) Cell viability of fibroblasts in the 3D myocardial tissue model. (H and I) The representative immunofluorescence staining against fibroblast activation markers including α-SMA and Vimentin in the 3D myocardial tissue model (H), with the quantification of fluorescent intensity (I). Scale bar, 100 μm. (J) The determination of fibrosis markers including COL1A1, COL3A1, SPP1, and Galectin3 in culture supernatant by ELISA assessments. Data are mean ± SEM. Statistical analysis was performed using one-way analysis of variance (ANOVA) among multiple groups, which was followed by an LSD post hoc test between two groups.

To investigate whether AH001 ameliorates fibrosis in the 3D myocardial tissue model through the RhoA-RhoGDIα axis (Figure 5A), we performed RhoGDIα knockdown in this model (Figure 7A). Quantification of Calcein-AM (a vital fluorescent marker for live cells) fluorescence intensity and stained area revealed that AH001 treatment significantly increased the Calcein-AM positive area, indicating enhanced cardiomyocyte viability (Figures 7B and 7C). However, the cardioprotective effects by AH001 were substantially attenuated upon RhoGDIα knockdown. Similarly, AH001-induced suppression of MRTFA nuclear translocation was abolished in RhoGDIα-knockdown conditions (Figures 7D and 7E). These results demonstrate that AH001 exerts its anti-fibrotic effect in a RhoGDIα-dependent manner, and stabilizes the RhoA-RhoGDIα complex to inhibit fibroblast proliferation and activation, promote cardiomyocyte survival, and block pro-fibrotic MRTFA signaling.

**Figure 7.**
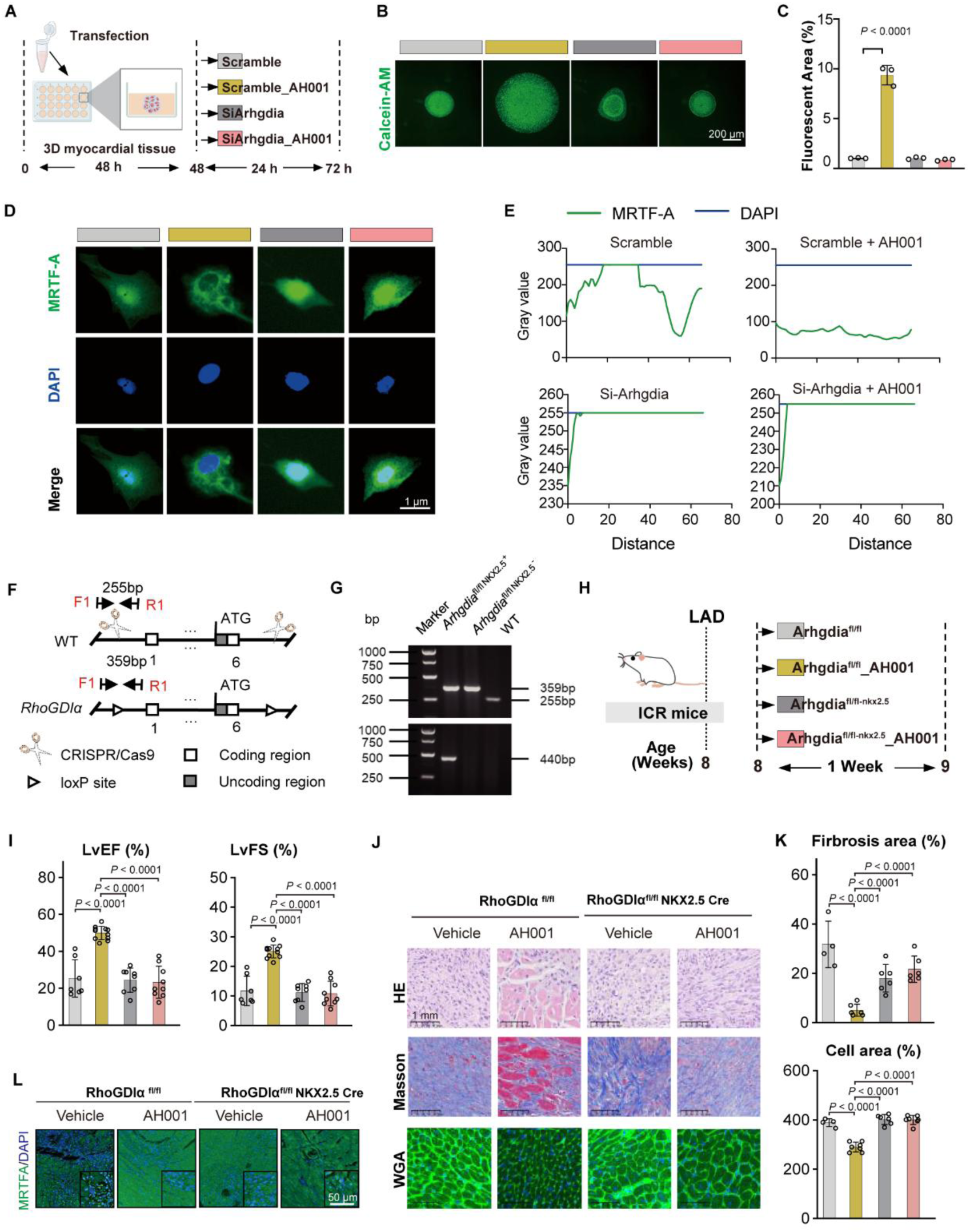
Knockdown of RhoGDIα in the 3D myocardial tissue model and in vivo. (A) The schema of knockdown RhoGDIα in the 3D myocardial tissue model, followed by the treatment of AH001 at 20 μM. (B and C) The representative images for the contractile phenotype of AC-induced fibrosis in the 3D myocardial tissue model (B), with fluorescent areas detected using Image J (C). Scale bar, 200 μm. (D and E) The representative images (D) of immunofluorescence staining against MRTFA, with quantification of nuclear MRTFA (E). Scale bar, 1 μm. (F and G) The schema of *RhoGDIα*^fl/fl^ NKX2.5 Cre mice construction (F) and genotyping of the *RhoGDIα*^fl/fl^ NKX2.5 Cre^+^, *RhoGDIα*^fl/fl^ NKX2.5 Cre^-^ and wildtype (WT) mice (G). (H) The schema of LAD mouse models construction with *RhoGDIα*^fl/fl^ and *RhoGDIα*^fl/fl^ NKX2.5 Cre mice, followed by the AH001 treatment at 10 mg/kg. (I) EF and FS in LAD mouse models described in (H) by the Echocardiographic assessment. (J and K) Representative images of Masson’s trichrome staining and WGA staining of a cross-sectional heart specimen, shown in the left ventricle region (J). Scale bar, 1 mm. Cardiac fibrosis was assessed by the fibrosis area and the cross-sectional area of cardiomyocytes (K). (L) Representative images of immunofluorescence staining against MRTFA on cross-sectional heart specimens, shown in the left ventricle region. Scale bar, 50 μm. Data are mean ± SEM. Statistical analysis was performed using one-way analysis of variance (ANOVA) among multiple groups, which was followed by an LSD post hoc test between two groups.

### AH001 inhibits the downstream signaling of RhoA activation in vivo

To validate the proposed functional mechanism of AH001 in vivo, we generated RhoGDIα^fl/fl^ mice using the CRISPR/Cas9-mediated genome editing, which were subsequently crossed with NKX2.5 Cre mice to generate myocardium-specific RhoGDIα knockout mice (RhoGDIα^fl/fl^ NKX2.5 Cre) (Figure 7F). After genotyping confirmation, mice were subjected to LAD surgery (Figures 7G-7H). Genetic ablation of RhoGDIα substantially attenuated the therapeutic effects of AH001, as evidenced by impaired recovery of cardiac function indicators, persistent fibrotic remodeling, unaltered myocardial hypertrophy, and no significant improvement in fibrosis areas or cardiomyocyte areas (Figures 7I-7K). Furthermore, genetic ablation of RhoGDIα abolished AH001-mediated suppression of MRTFA nuclear translocation, as evidenced by no significant intergroup difference of the fluorescence signal of nuclear MRTFA in myocardial tissue sections (Figure 7L). These results conclusively demonstrate that AH001 exerts its cardioprotective effects through the RhoA-RhoGDIα axis, effectively inhibiting downstream RhoA activation signaling in vivo.

## DISCUSSION

As RhoA controls cytoskeletal reorganization and cell migration—processes central to myocardial hypertrophy and fibrosis—its aberrant activation contributes to myocardial remodeling pathogenesis. Consequently, inhibiting RhoA activation has emerged as a promising therapeutic strategy to attenuate myocardial remodeling and delay HF progression. However, RhoA has long been considered challenging as a drug target. In this end, we proposed the drug screening strategy targeting RhoA via a cryptic pocket near GDP to constrain RhoA in the GDP-bound state. A nature product, AH001, was identified to occupy the pocket and interact with RhoA and GDP. We established multi-species HF models incorporating zebrafish, mouse, and rat, combined with the 3D myocardial tissues to simulate HF of diverse etiologies. We demonstrated that AH001 effectively improved multiple cardiac function indicators and inhibited myocardial hypertrophy and fibrosis in these models. These therapeutic effects of AH001 resulted from the maintenance of RhoA-RhoGDIα complex and the inhibition of MRTFA nuclear translocation, which was further validated by knockdown of RhoGDIα in the 3D myocardial tissue model and the myocardium-specific RhoGDIα knockout mouse model. Additionally, we observed elevated RhoA activation in fibroblasts and cardiomyocytes from HF patients of diverse etiologies by the multi-omics data analysis. We demonstrated impaired RhoA-RhoGDIα interaction in myocardium from HF patients, evidenced by reduced spatial colocalization in immunofluorescence analysis of tissue sections. Our work provides new insights into drug screening and design targeting previously considered “undruggable” GTPases such as RhoA. Our work also demonstrated that inactivation of RhoA blocked the pathological process of myocardial remodeling, offering a previously unrecognized therapeutic strategy for myocardial remodeling, and thereby targeting HF.

The Rho GTPases, including RhoA, are crucial cellular regulators and control various physiological processes underlying cytoskeletal organization and cell motility.^25, 26^ Rho GTPases have long been deemed “undruggable”, due to two major challenges: (i) the strong binding to endogenous substrates of GTP or GDP, and (ii) the absence of well-defined binding pockets.^27^ Addressing the second challenge, targetable switch II pockets were identified by the systematic survey across members of various GTPases.^17, 28, 29^ Equivalent cysteine mutations were introduced into other GTPases based on sequence alignment with K-Ras.^17^ Cysteine mutated GTPases undergo the covalent modification by optimized K-Ras (G12C) inhibitors binding in the switch II pocket, demonstrating the conservation of switch II pockets. However, only RabL5 (in the Ras superfamily) has a native cysteine at the mutation position. Thus, it remains unknown whether this pocket is targetable for wild-type proteins. Recently, a novel allosteric pocket with the conserved cysteine 107 was identified by molecular dynamics simulation.^20^ The inhibitor DC-Rhoin formed the covalent modification to cysteine 107 and inhibited activation of RhoA. However, it is shown that the cysteine 107 can be natively mutated into a serine or tyrosine at this position in the Ensembl database (Variant ID: rs2107830328), which will lead to decreased effects of the covalent inhibitor on mutant proteins. Moreover, the first challenge remains unresolved.

In this study, we discovered a cryptic pocket located near GDP from the structural difference analysis on RhoA-GTP and RhoA-GDP, and identified AH001 bound in the cryptic pocket via hydrophobic interactions with T37 and Q63 of RhoA. AH001 also forms a hydrogen bond with GDP, thereby it restricting RhoA in the GDP-bound state. Crucially, unlike sustained RhoA inhibition, which may lead to a shorter lifespan and the reduced ejection fraction seen in cardiomyocyte-specific RhoA knockout mice,^30^ reversible inhibitors maintain physiological pathway dynamics, because the reversible binding enables concentration-dependent activity modulation (an ‘on-off’ switch effect), thereby reducing overdose risks and avoiding off-target effects caused by covalent modification.^31, 32^ We address the two aforementioned challenges for undruggable RhoA, and also provide opportunities for the future development of reversible inhibitors for various GTPases.

Activated RhoA-GTP allows F-actin polymerization and depletes cytoplasmic G-actin to release MRTFA.^16^ Free MRTFA enters the nucleus to initiate SRF-dependent gene transcription, driving pro-fibrotic and hypertrophic gene expression.^13, 16^ While RhoA overexpressing causes lethal HF in mice,^14^ whether the progression of HF specifically requires MRTFA dysregulation such as persistent nuclear accumulation or aberrant transcriptional upregulation is unknown. Additionally, although RhoGDIα canonically sequesters RhoA-GDP in the cytoplasm to inhibit RhoA activity,^33^ whether the RhoGDIα-mediated fluctuations in RhoA activity may feedback to control MRTFA nuclear translocation or expression levels remains unexplored.

In this study, we demonstrated that AH001 facilitated the binding of RhoA-RhoGDIα to constrain RhoA in the GDP-bound state, suppressing MRTFA nuclear translocation to reduce pro-fibrotic and hypertrophic gene expression. Additionally, we identified a *Mrtfa*+ cardiomyocyte subpopulation through the scRNA-seq analysis, which can interact with fibroblasts mutually through the paracrine SPP1 signaling. AH001 effectively reduced the cell-cell interaction between *Mrtfa*+ cardiomyocytes and fibroblasts, with downregulation of *Spp1* in *Mrtfa*+ cardiomyocytes and fibroblasts and disease-associated fibroblast markers such as *Postn*, *Acta2*, and *Runx1*. Consistently, *Mrtfa* knockout mice exhibited diminished fibroblast activation and decelerated fibrosis progression.^21, 34^ Specifically, in the 3D myocardial tissue model simulating cell-cell interactions between *Mrtfa*+ cardiomyocytes and fibroblasts, we demonstrated that AH001 suppressed proliferation and activation of fibroblasts, alleviated the survival of cardiomyocytes, and decreased the secreted SPP1 in the culture supernatant in a dose-dependent manner. Additionally, the knockdown of RhoGDIα in the 3D myocardial tissue model reversed AH001-mediated suppression of MRTFA nuclear translocation, which again proved that AH001 acted on the RhoA-RhoGDIα-MRTFA axis in both fibroblasts and cardiomyocytes. Our findings not only establish MRTFA+ cardiomyocytes as active contributors in fibrotic signaling, but also suggest SPP1 as a viable preclinical biomarker for potential clinical translation of AH001.

The intrinsic difficulty in pharmacologically modulating RhoA has driven the development of alternative strategies to inhibit the RhoA signaling pathway. Rhosin (G04) can bind to RhoA (K_d_ = 354 ± 48 nM) and disrupt the binding between RhoA and its guanine nucleotide exchange factors (GEFs) to suppress platelet activity.^35, 36^ However, the pharmacokinetic profile of Rhosin and its precise molecular mechanism remain uncharacterized. CCG-1423 with its derivative CCG-203971 inhibits MRTFA/SRF-dependent transcriptional activation (IC_50_ = 0.64 μM) to exhibit antifibrotic effects, with the direct molecular target is unclear.^37^ Although pan-ROCK (Rho-associated coiled-coil-containing kinases) inhibitors such as Fasudil and Ripasudil ameliorate cardiac fibrosis in murine models,^38, 39^ emerging evidence strongly implicates that these effects are primarily mediated through ROCK2 inhibition while ROCK1 is not necessary for causing myocardial hypertrophy.^40^ The non-selectivity of clinically available ROCK inhibitors may thus compromise therapeutic efficacy and increase risks of potential adverse effects. Unlike Rhosin and CCG-1423, we discovered AH001 directly bound to RhoA-GDP and stabilized the RhoA-RhoGDIα complex without disrupting physiological RhoA cycling in this study. By contrast to pan-ROCK inhibitors that may compromise therapeutic efficacy on myocardial hypertrophy, AH001 alleviated both myocardial hypertrophy and fibrosis via suppressing MRTFA nuclear translocation in multiple HF models. Collectively, AH001 represents a mechanistically distinct and therapeutically superior RhoA-targeting inhibitor by overcoming long-standing barriers in this field.

In conclusion, we have proposed a drug screening strategy targeting undruggable RhoA and established the RhoA-RhoGDIα-MRTFA axis that controls myocardial remodeling including hypertrophy and fibrosis. Our strategy to identify compounds bound in the pocket near GDP offers opportunities for developing reversible inhibitors targeting various GTPases. Our multi-omics analysis confirmed the clinical relevance of the RhoA-RhoGDIα-MRTFA axis in the HF patients, while integrated cross-scale experimental platforms (in vitro primary cells, 3D myocardial tissues, and in vivo) demonstrated consistent therapeutic effects of inhibiting RhoA on myocardial remodeling. This work provides not only therapeutic potential but also a generalizable platform for developing HF treatments.

## ACKNOWLEDGMENTS

We are sincerely grateful to Prof.Yinchu Shen for his support in the help for this project. We wish to thank Profs. Jingyan Zhang, Yan Zhang, Zijian Li, and Guoping Shi, for technical advice, suggestions, and critical reading of the manuscript. This work was supported by the National Natural Science Foundation of China (82425104 to H.L., 82222075 to C.L.) and the National Key Research and Development Program of China (2022YFC3400501 to H.L.), the Science and Technology Commission of Shanghai Municipality (24JS2830200), and the Shanghai Municipal Education Commission (2024AI01014).

## AUTHOR CONTRIBUTIONS

H.L. initiated the project. H.L., C.L., J.Z., Y.W., and H.T. supervised the overall project. M.X., Y.L., Z.Y., Y.Z.W., and X.L. wrote the original manuscript and prepared the figures. M.X. performed the CITE-seq, snRNA-seq, and scRNA-seq analysis. Y.L. performed the in vitro cellular experiments with help of P.X., T.Z., Y.W., and J.W.W. H.L., Z.Y., and S.R. performed the virtual screening and molecular dynamic simulation to discover the lead compound. X.L. constructed the 3D myocardial tissue model. J.L. and C.L. performed the SPR assay. L.Z., X.P., and L.C. purified protein and solved the X-ray complex crystal structure. Y.W., C.L., H.W., and X.S. performed the LAD model in mice and rats. H.H., J.C., and H.T. conducted the cardiac hypertrophy mouse model. Y.Z, and X.L. participated in the zebrafish model. Y.L and Y.D conducted Doxorubicin-induced mouse models. Y.Z.W. participated in the preparation of transgenic mice. H.J. and W.M., T.Y., S.T., Z.J.Z. detected the target of AH001. J.W., Z.Z., and C.L. performed the detection of RhoA-GTP/GDP ratio. H.L., C.L., R.W., Y.D., and Y.L. provided suggestions and M.X. edited the manuscript; All authors read and approved the final manuscript.

## STAR METHODS

### Animals

The ICR mouse and the SD rats for the Left anterior descending coronary artery ligation model were bought from SPF (Beijing) Biotechnology Co., Ltd. C57BL/6J mice for Ang II induced models and Dox-induced models were purchased from Cyagen Biosciences Inc and GemPharmatech LLC. The *RhoGDIα*^fl/fl^NKX2.5 Cre mice were on a C57BL/6J background and provided by Cyagen Biosciences Inc. All mice were housed under standard temperature (22 ± 2°C) and humidity (60 ± 10%), with a 12 h light-dark cycle.

### Ethics statements

Animal studies were carried out in compliance with the recommendations in the Guidelines on the Care and Use of Animals for Scientific Purposes of the National Advisory Committee for Laboratory Animal Research. All experimental procedures were approved (HFLIT 20220713001, BUCM-2023052501-2119, BUCM-2023050902-2078, SHOU-dw-2021-042) skillful experimenter were responsible for all animal experiment procedures according to the laws governing animal research in China. The pathological sections from healthy donors and heart failure patients used in this study were de-identified, and the research procedures complied with the ethical principles of the Declaration of Helsinki and China’s Ethical Review Measures for Biomedical Research Involving Human Subjects (KS2023083).

### Isopropyl hyperglandrin-induced myocardial hypertrophy model in zebrafish

The zebrafish myocardial hypertrophy model by Isopropyl hyperglandrin (ISO). After the initial sample concentration test, the samples were divided into 4 groups: WT group, model group, LCZ696-treated Group and AH001-treated Group. Cardiac function was measured 72 h after drug administration, and ejection fraction and other related indexes were counted.

### Left anterior descending coronary artery ligation model

Permanent left anterior descending (LAD) coronary artery ligation was performed as previously described with minor modifications. ICR mice or SD rats at the age of 8-10 weeks, were anaesthetized using pentobarbital sodium at 50 mg/kg body weight by intraperitoneal injection, and then intubated and ventilated with air (supplemented with oxygen) using a small-animal respirator. A thoracotomy was performed in the fourth left intercostal space. The left ventricle was visualized and the pericardial sac was ruptured to expose the LAD coronary artery. The LAD was permanently ligated using a 7-0 Prolene suture. The suture was passed approximately 2 mm below the tip of the left auricle. Marked colour changes of the ischaemic area and ECG changes were monitored as an indication of successful coronary artery occlusion. The thoracotomy was closed with 6-0 Prolene sutures. Sham-operated mice underwent the same procedure without coronary artery ligation. The endotracheal tube was removed once spontaneous respiration resumed, and the mice were placed in a warm recovery cage maintained at 37 °C until they were completely awake. At the indicated time points, the mice were euthanasia and the tissues were subsequently collected for analysis.

### Ang II induce mouse models

C57BL/6J mice (8 week) were subcutaneously implanted with osmotic minipumps to deliver angiotensin II (Ang II) at a dose of 3 mg/kg/day. The pumps were surgically inserted under anesthesia by Isoflurane (2%), and Ang II infusion was maintained for 28 days. Starting from the third week post-pump implantation, mice received either AH001 (10 mg/kg/day) or LCZ696 (66.7 mg/kg/day) vehicle via oral gavage for 28 days. Cardiac function and blood pressure was assessed using an echocardiograph (Vevo TM2100, Visual Sonics). Tissue samples (heart) were collected for histopathological analysis (Masson’s trichrome, and WGA staining) to evaluate cardiac remodeling and fibrosis.

### Dox-induced mouse models

Following a 1-week adaptive feeding period for 6-week-old male mice, 8 mice were injected with normal saline as the control group, and the remaining 50 mice were intraperitoneally injected with doxorubicin (Dox, purchased from MCE, Shanghai, China). The total cumulative dose of doxorubicin was 20 mg/kg, administered weekly at a dose of 5 mg/kg, with injections given at the same time daily for 4 consecutive weeks. After the fourth Dox injection, cardiac function was evaluated using a Digital Color Doppler Ultrasound System. Mice with a left ventricular ejection fraction (EF) of approximately 50% were considered to have successfully established the heart failure model. Subsequently, these model mice were randomly assigned into five groups (n=10/group) for intragastric administration: Dox group, daily administration of saline; LCZ696 group, daily administration of 60 mg/kg LCZ696; AH001 low/mid/high dose groups, daily administration of 1 mg/kg, 10 mg/kg, and 50 mg/kg AH001, respectively. Mice in the control group were intragastrically administered with an equal volume of normal saline. Following 12 days of treatment, echocardiography was repeated, and then the mice were sacrificed.

### Echocardiographic assessment

The ICR mice and SD rats were anesthetized by pentobarbital sodium. Cardiac function was assessed using an echocardiograph (Vevo TM2100, Visual Sonics). The main parameters included the left ventricular ejection fraction (LVEF), short-axis shortening fraction (LVFS), left ventricular end-diastolic internal diameter (LVID; d), and left ventricular end-systolic internal diameter (LVID; s).

### Genotyping

The mice were screened by PCR using genomic DNA extracted from mouse tails.

For qualitative PCR, two primer pairs for *Arhgdia* were used as follows.

F1: TCTTGGTAGTTTCAGCCATTCGTGT,
R1: TACCACCCGGTGAAATCTGGAAATT;
F2: AACAGGAAACCGAGCCTCTGCA,
R2: TTTAGACTGACTCCTGTAAAGGCAGG;

The PCR reactions in 20 µL volumes were carried out using 10 ng genomic DNA, OneTaq® 2 × Master Mix (Ipswich, New England Biolabs), and 10 pM of oligonucleotide primers. The reaction mixture was incubated at 94 °C for 3 min, followed by 32 cycles at 94 °C for 30 s, at 57.5 °C for 30 s, and at 68 °C for 25 s, and a final cycle at 68 °C for 5 min in CFX96TM Real-Time System (Hercules, Bio-Rad). Agarose gel electrophoresis was used to separate the PCR products, which were visualized using ChemiDocTM MP Imaging System (Bio-Rad).

### Histology assays

Mice and Rats were euthanized at different time intervals. The hearts were collected and sliced into frozen sections that were 5 μm thick. To determine the histologic characteristics of myocardial infarction, Masson trichrome staining was performed according to the instructions provided by the manufacturer. The infarct size was determined by calculating the ratio of collagen (blue) to myocardial muscle (red) in the frozen heart section. To assess tissue cell area, 5 μm thick sections of the heart tissues were embedded in paraffin and mounted on poly lysine glass slides. The slices were then H&E and WGA stained. Image J software was used to analyze the staining data.

### The pharmacokinetic property determination of AH001

Healthy adult SD rats (male, body weight 180-220 g) were acclimatized for 1 week prior to the experiment with free access to food and water, under a 12-hour light/dark cycle. SD rats were randomly divided into two groups (n=3): intravenous (IV) or intragastric (IG) administration with the same dose of 10 mg/kg. Blood was collected via the orbital venous plexus in rats at predosing (0 min) and at various time points postdosing in the IV and IG groups (5 min, 15 min, 30 min, 1 h, 2 h, 4 h, 8 h, 12 h, 24 h). After centrifugation at 15000 rpm for 10 min at 4 °C, the blood samples were analyzed using an AB SCIEX Qtrap 5500 mass spectrometry system, and pharmacokinetic parameters were derived using WinNonLin 8.0 for data analysis. Chromatographic separation was achieved using an ACQUITY UPLC® BEH C18 column (2.1 × 50 mm, 1.7 μm particle size) under optimized conditions.

### The GLP-compliant toxicity studies of AH001

The safety pharmacology studies for AH001 were conducted by the Center for Drug Safety Evaluation, Chinese Academy of Sciences, Shanghai Institute of Materia Medica, which holds GLP certification by National Medical Products Administration (NMPA) and ISO 9001:2000 Quality Management System certification.

### Isolation of ventricular cardiomyocytes and fibroblasts from neonatal rats

The neonatal rats used were brought from Dachengyinuo Biomedicine (Shanghai, China) Co., Ltd. Wistar rats were handled in accordance with the approved license (HFLIT 20220713001). Neonatal rat ventricular cardiomyocytes were isolated from 2-day-old (P2) male wild-type Wistar rats. Hearts of rats were pulled out by forceps and transferred into a 60-mm Petri dish containing 1 × phosphate-buffered saline (PBS). The remaining blood was pumped out of the hearts. Both atria and the connective tissue were removed. For each preparation, a batch of hearts were cut into small pieces, and the tissues were digested with 5 mL 0.25% Trypsin-EDTA overnight at 4℃. Then, 0.25% Trypsin-EDTA was removed followed by second digestion with Dulbecco’s modified Eagle medium and Ham’s F12 medium (PM150312A, Procell) containing 0.1% collagenase II (LS004177, Worthington) and 1% Bovine serum albumin (S25762, Yuanye) was filtered with 0.22 μm syringe filter (BS-PES-22, Biosharp).

Briefly, adding an appropriate amount of sterilized PBS to overnight digestion, washing it three times, and then adding 1 mL of the digestive solution. Shaking gently in a 37°C constant temperature water bath for 1 min. Adding 7 mL of digestive solution mentioned above, gently shaking in a 37 °C water bath for 1.5 min, and collecting 7 mL cell digestive solution into an RNA/DNA-Free centrifuge tube (collect three times) with 21 mL cardiomyocyte medium. This medium consists of Dulbecco’s modified Eagle medium and Ham’s F12 medium (PM150312A, Procell) containing 10% fetal bovine serum (10091148, Gibco) and 1% penicillin-streptomycin (15140122, Gibco). Centrifuging at 300 g for 5 min.

Cell suspensions containing cardiomyocytes and cardiac fibroblasts, isolated from whole heart extracts, were seeded into 75 cm² polystyrene culture flasks, as previously reported.^41^ The cells were resuspended in DMEM supplemented with 10% fetal bovine serum (FBS) and 1% penicillin-streptomycin, then incubated at 37°C under 5% CO₂. After approximately 2 hours of differential adhesion, cardiac fibroblasts adhered to the flask surface, while cardiomyocytes remained in suspension due to their inability to attach to uncoated plastic. The non-adherent cell suspension (enriched with cardiomyocytes) was carefully transferred to a new vessel for further processing.

### Cell culture

Rat H9C2 cells was purchased from ATCC (Manassas, VA, USA) and maintained in DMEM with 10% FBS and 1% penicillin-streptomycin, at 37 °C in a humidified incubator containing 5% CO_2_.

Hypoxia condition, H9C2 cells were exposed to hypoxia for 6 hours as described previously (Fan S.Y., et al. Quantum sensing of free radical generation in mitochondria of single heart muscle cells during hypoxia and reoxygenation, ACS Nano, 2024). We used an AnaeroPack system (Mitsubishi, Tokyo, Japan) to offer an anaerobic condition (AC). Briefly, DMEM without glucose and corresponding treatments were added to the cells, which were then placed into an anaerobic bag and cultured at 37°C for 6 hours.

### Cell migration assays

Migration of CFs was tested by Transwell and scratch assay. In the Transwell assay, CFs at a density of 1×10^5^ cells per well were seeded into the upper chamber (12 μm pore size; LABSELECT, Hefei, China) in 100 μL serum-free DMEM, 500 μL DMEM medium with or without TGF-β1 (10ng/mL) was added in the lower chamber. Alternatively, culture medium for H9C2 cells anaerobic or aerobic conditions was introduced. After 24 hours’ incubation, cells were fixed with 4% paraformaldehyde, stained with crystal violet (Beyotime Biotechnology, Shanghai, China), and photographed. For scratch assay, a sterile 200-μL pipette tip was used to create a uniform scratch across the CFs monolayer in 6-well plates. Cells were then incubated in serum-free DMEM with or without TGF-β1 for 24 hours to allow migration into the wounded area. Four random fields per well were imaged at 0- and 24-hours post-scratching. Wound closure was quantified by measuring the mean linear movement of the wound edges over 24 hours, calculated as the relative migration speed.

### Knockdown RhoGDIα in cardiomyocytes and fibroblasts

To knock down RhoGDIα, small interfering (si)RhoGDIα was transduced into H9C2 or fibroblasts. The sequences were 5′-GGUGUGGAGUACCGGAUAA-3′ and 3′- UUAUCCGGUACUCCACACC-5′ for rat *Arhgdia*. The knockdown efficiency was determined by western blotting.

Seed primary cardiac fibroblasts and cardiomyocytes in 6-well plates and culture until 70% confluency. Dilute Lipofectamine RNAiMAX in Opti-MEM. Mix with siGDI in Opti-MEM, incubate 5 min at RT. Add the complex dropwise to cells. Gently swirl plates. Incubate at 37°C or 48 h. Collect the transfected cells and culture with collagen to establish the 3D myocardial tissue model.

### The Proximity ligation assay (PLA) assay

PLA, widely used for detecting in situ protein–protein interaction was used to detect the in situ interaction of endogenous RhoA and RhoGDI proteins in cells by employing Duolink secondary antibodies and a detection kit (DUO92101, Sigma) according to the manufacturer’s instructions. In brief, 1 × 10^5^ cells were seeded in 24-well plate, in a cell incubator at 37℃ overnight. The plasmid was transferred in cells for 48h. Then the cells were fixed with 4% paraformaldehyde at room temperature for 10 min, washed with 1 × PBS for 5 min, and repeated 3 times. Cells were incubated with anti-RhoA (1:200 in TBST, A0272, Abclonal) and mouse anti-flag (1:500 in TBST, AE005, Abclonal) antibodies together with Duolink secondary antibodies. Ligation steps were performed to ligate the secondary antibodies that are in close proximity. 4′,6-diamidino-2-phenylindole (DAPI) was used for counterstaining the nucleus. After standing for 15 minutes and images were taken under confocal microscope.

### Western blotting

Cells were lysed with lysis buffer containing protease inhibitor cocktail (CW2200S, CWBIO). Protein concentrations were determined by the BCA Protein Assay Kit (P0010, Beyotime). Equivalent levels of protein samples were electrophoresed by loaded onto 10% SDS-PAGE and transferred to polyvinylidene difluoride membranes. The membranes were blotted with 5% non-fatty milk in tris-buffered saline Tween-20 (TBST) for 2 h at room temperature and incubated with primary antibodies overnight at 4°C. Primary antibodies were diluted as follows: anti-RhoA (1:1000 in TBST, 21017, NewEast), anti-RhoGDI (1:1000, Abcam, ab133248, England), Anti-RhoA-GTP (1:1000 in TBST, 26904, NewEast), anti-Collagen I (1:1000 in TBST, ab270993, Abcam), HRP-conjugated anti-α-Tubulin antibody (1:10000 in TBST, HRP-66031, Proteintech), anti-COL3(1:2000 in TBST, A0817, abclone, USA), anti-VIM(1:1000 in TBST, YP-Ab-03212, UpingBio, China), anti-SRF(1:1000 in TBST, YP-Ab-02042, UpingBio, China), anti-FN1(1:1000 in TBST, YP-Ab-17113, UpingBio, China), anti-CFN(1:1000 in TBST, YP-Ab-03106, UpingBio, China), p-CFN(1:1000 in TBST, YP-Ab-03011, UpingBio, China), HSP90 (1:10000 in TBST, 13171-1-AP, Proteintech: Chicago, USA), ANP (1:2000 in TBST, DF6497, Affinity), BNP (1:2000 in TBST, DF6902, Affinity), and MYH7 (1:2000 in TBST, DF12122, Affinity).

### ELISA for indicator proteins determination in culture supernatant

After H9C2 cells were cultured in glucose-free DMEM under anaerobic or aerobic conditions for 6 hours, the culture medium was collected to detect the fibrogenic secretory factors. ELISA kits were performed to test the level of SPP1(UpingBio, SYP-R1889, Hangzhou, China), CTGF (UpingBio, SYP-R0079, Hangzhou, China), TGF-β1(ZciBio, CSB-E08393r, Shanghai, China) according to the procedures described in the instruction manual.

Collect supernatant samples from the 3D myocardial tissue model cultures. The samples were centrifugated at 3000×g for 10 min at 4°C. After centrifugation, the supernatant was detected for enzyme-linked immunosorbent assays of mouse collagen type I alpha 1 (COL1a1) ELISA kit (Bioswamp, China), mouse collagen type III alpha 1 (COL3a1) ELISA kit (Bioswamp, China), mouse osteopontin (OPN/SPP1) ELISA kit (Bioswamp, China), and mouse galectin-3 ELISA kit (Bioswamp, China).

### Live staining of the 3D myocardial tissue model

For live staining, cells were washed twice with PBS and then incubated with the dye mixture (2 mL calcein-AM in 1 mL PBS) for 10 min at 37°C. Subsequently, the living cells and apoptotic cells were observed and imaged using a confocal microscope.

### Immunofluorescence staining of cells or mouse heart tissue sections

Cells on coverslips or tissue sections were fixed and permeabilized, blocked by 2% bull serum albumin for 1.5 h. Then, the coverslips or tissue sections were incubated with anti-MRTF-A antibodies (1:200, Proteintech, 21166) overnight at 4 °C. Following three rinses with PBST (PBS containing 0.1% Tween, 5 minutes per rinse), the coverslips or sections were incubated with Alexa Fluor 555-conjugated secondary antibody (1:500 dilution, Abcam, ab150118) for 1 hour at room temperature in the dark. After washing cells or tissue sections with PBST, they were incubated with DAPI for 10 minutes at room temperature in the dark. Following 3 additional PBST washes, cells or tissue sections were mounted using antifade mountant (Servicebio). Confocal laser scanning microscopy (Nikon) was used to capture images, and fluorescence intensity quantification was performed using ImageJ software.

For F-actin determination, cells on coverslips were fixed and permeabilized as above. The cells were stained with actin-tracker red rhodamine (Beyotime, C2207) at room temperature (RT) for 60 minutes, followed by three rinses with PBST. Followed by DAPI incubating for 10 minutes at RT in the dark, an additional 3 times PBST washes. The coverslips were subsequently mounted, and confocal microscopy images were captured using a Leica confocal laser scanning microscope.

### Immunofluorescence staining in the 3D myocardial tissue model

The cells were fixed with 4% paraformaldehyde (PFA) for 15 min at room temperature (RT), followed by three washes with phosphate-buffered saline (PBS). Permeabilization was performed using 0.5% Triton X-100 for 20 min at RT. After washing with PBS, nonspecific binding sites were blocked with 1% bovine serum albumin (BSA) at 37°C for 1 h in a humidified incubator. Primary antibodies were applied and incubated overnight at 4°C. Subsequently, cells were washed with PBS and incubated with species-matched fluorescent-conjugated secondary antibodies for 1 h at RT in the dark. Nuclear counterstaining was achieved with DAPI for 5 min. Following final PBS washes, fluorescent signals were visualized and imaged using a laser scanning confocal microscope (Leica Wetzlar, Germany).

The antibodies and dyes used were as follows: anti-alpha-actinin (α-actinin) antibody (Proteintech, USA, 1:250), anti-vimentin antibody (Origene, USA, 1:250), anti-alpha smooth muscle (α-SMA) antibody(Abcam, England, 1:250), DAPI (Abcam, 1:200), Goat anti-rabbit IgG (Cohesion Biosciences, England, 1:200), and Goat anti-chicken IgG (Cohesion Biosciences, 1:200).

### Multiplex immunohistochemistry (mIHC) with tyramide signal amplification (TSA) cyclic staining

Paraffin-embedded sections from healthy donors, heart failure patient, and the 3D myocardial tissue model samples were subjected to heat-induced antigen retrieval using citrate buffer and rinsed three times with PBS. Endogenous peroxidase activity was quenched by incubating slides with 3% hydrogen peroxide for 15 min at RT. Primary antibodies were applied and incubated overnight at 4°C. HRP-conjugated secondary antibodies were incubated for 1 h at 37°C, followed by tyramide-fluorophore conjugates for 10 min. Slides were imaged using an Olympus SpinSR10 confocal microscope (Olympus, Japan) with predefined exposure settings. Antibody complexes were removed through microwave treatment for 15 min. This cycle was repeated for subsequent targets. After completing all iterative rounds, nuclear counterstaining was performed with DAPI for 5 min. Final high-resolution z-stack images were acquired using the Olympus SpinSR10.

The antibodies and dyes used were as follows: anti RhoA rabbit polyclonal antibody, (NewEast Biosciences, USA, 1:250), anti-RhoGDI (Abcam, 1:250), and IRISKit® HyperView mIF Kit.

### Cell viability assay

In brief, 10% CCK-8 in DMEM was added to the wells and incubated at 37°C for 2 h. The optical density of the cells in each well was then measured using a microplate reader (PerkinElmer, Waltham, MA, USA) set to a wavelength of 450 nm.

### RhoA pull-down activation assay

RhoA activity was assessed using the RhoA Pull-Down Activation Assay Biochem Kit (Cytoskeleton, BK036) following the product protocols. Briefly, proteins were rapidly extracted from cells, with 50 μL reserved for western blot analysis to quantify total RhoA levels. The active RhoA fraction (RhoA-GTP) was then captured by rhotekin-RBD beads through gentle agitation at 4 °C for 1 hour. After centrifugation at 5,000 × g for 1 minute at 4 °C, the beads were washed once with buffer, resuspended in 20 μL of 2× reducing SDS-PAGE sample buffer, boiled for 5 minutes, and subjected to western blotting.

### RhoA activity detection

1×10^6^ cells were seeded and stimulated by Ang II. The cells were washed with PBS twice, quickly joined 500 edged up protein cracking liquid containing protease phosphatase, scrape cells using a scraper, ice cracking 15 min, 4℃ under the condition of 12000 rpm centrifuge for 10 min. To proceed affinity precipitation, 1 ng of RhoA-GTP antibody was added and protein A/G agarose was thoroughly mixed, 20 ml were added and incubated at 4℃, centrifuged at 500 rpm for 10 s to remove the supernatant, washed 3 times, and finally 20 ml of protein loading buffer was added at 100℃ for 5 min. Then, the samples were proceeded with the test of RhoA activity in western blotting assay.

### RhoA GTP/GDP ratio

The GRI SerHR-RhoA-GTP and GRI SerHR plasmids in this experiment were provided by Wang Jing’s research group of Peking University. The specific operation flow of the experimental steps is as follows: 1 × 10^4^ cells were seeded in 96-well plates (P96-1-N, Cellvis). The transfection reagent prepared according to the instructions of lipo3000 reagent (L3000015, Thermo Fisher), add opti-MEM (1985062, Gibco) to prepare a suitable volume of transfection reagent mixture. Leave the mixture at room temperature for 15 min, remove the cell culture medium from the cell culture plate, and wash it once with phosphate-buffered saline (PBS). The cells were added into the plasmid mixture after resting, and the cells were placed back in the incubator for further culture. After 6 h, the cell medium containing the mixed solution was removed, and the complete medium was added for further culture for 18 h, and then AH001 was added for 12 h. The cell intensities excited at 488 nm and 405 nm (F488/F405) upon to GTP was detected in confocal mode, and the fluorescence intensity ratio was calculated by image J software (version 1.54 f) as RhoA GTP/GDP conversion rate.

### Surface Plasmon Resonance (SPR) protein molecular interaction analysis experiment

The specific operation process of Surface Plasmon Resonance (SPR) for the verification of AH001 and protein was shown as follows: HEK 293F cells were transfected by full-length of RhoA cDNA with His tag for 48h, RhoA protein was extracted with protein cracking solution and purified. SPR on-board preparation was operated. A new BIAcore CMP-5 dextran chip was inserted and PBS was used as the working fluid to balance the system. The optimal buffer conditions and concentration were determined experimentally to minimize the target and ligand proteins non-specifically bound to the carboxylate esterification dextran. Dissociation constant determination: change the contact time and protein concentration, and then detect the separation of the target protein from each ligand density. The rate of separation should be constant and independent of the concentration of the surface density of the ligand. The ligand density should be optimized to obtain sufficient molecular surfaces that present a clear binding reaction. And dilute sufficiently to measure the dissociation rate constant that should be constant. The separation rate and binding rate constants were calculated using BIA evaluation software point-and-click. The equilibrium constant was calculated by averaging the binding and separation rates of the three target protein concentrations.

### Molecular dynamics (MD) simulations and virtual screening

We performed three independent molecular dynamics simulations for both the RhoA^WT^- GDP (PDB code: 1FTN) and RhoA^G14V^-GTPγS (PDB code: 1A2B). All simulations were performed using the PMEMD module of AMBER20 software package. These systems were solvated in a TIP3P water box of 10 Å, employing the AMBER14SB force field. Energy minimization was accomplished using the steepest descent algorithm for 5000 steps, followed by an additional 5000 steps using the conjugate gradient algorithm. Then, the systems were heated up progressively from 0 to 300 K for 200 ps using the Langevin thermostat in NVT ensemble. A restrained 200 ps production run was performed under 1.0 atm pressure with the temperature keeping constant at 300 K using Berendsen barostat in NPT ensemble. Finally, after the removal of position restraints, a production run was performed for 600 ns with a 2-fs step.

After performing clustering analysis on the MD trajectory of the RhoA^WT^-GDP system, a representative structure was selected for virtual screening. The protein structure was prepared using the Protein Preparation Wizard in Maestro (Schrödinger Inc, version 10.1). TOPSCIENCE natural product library and in-house library (∼180,000 compounds) were prepared and filtered using the LigPrep module. We carried out Glide HTVS, Glide SP, and Glide XP, and retained the top 20%, 20% and 20% of ligands for each stage, respectively. The top-ranked 1,019 molecules were retained for further visual observation. Taking into consideration ligand binding poses, structural diversity, and novelty, 100 candidates were selected from the pool of 1,019 compounds and subsequently subjected to activity testing.

### Crystallization and structure determination

The purified RhoA used for crystallization was concentrated to more than 15 mg/mL. Co-crystallization of RhoA with AH001 was performed at 16℃ using the vapor diffusion method.^42^ X-ray diffraction data were collected at 100 K on beam line BL02U1 at the Shanghai Synchrotron Radiation Facility (SSRF). The raw data were processed using the MOSFLM and SCALA programs from the CCP4 suite.^43^ The crystal structure of RhoA bound with AH001 was determined by molecular replacement using PDB entry 1FTN (without ligands and water molecules) as the search model. Structure refinement was conducted with REFMAC5. After cycles of refinement and model building with COOT.^44^ Statistics of data collection, processing and refinement are listed in Table S1. The molecular graphics were prepared with PYMOL (http://www.pymol.org) to generate the figures.

### The scRNA-seq library preparation on mouse heart specimens

Whole heart specimens were obtained from LAD mouse models and were dissociated to prepare single-cell suspensions. Single-cell suspensions were loaded onto the 10x Genomics Chromium Controller using the Chromium Single Cell 3′ Reagent Kit (PN-1000075, 10x Genomics, Pleasanton, CA, USA) following the manufacturer’s protocol. Briefly, viable cells (≥85% viability) were resuspended in PBS + 0.04% BSA at a target concentration of 700-1,200 cells/μL. Cells were combined with reverse transcription (RT) master mix and partitioning oil, then emulsified into Gel Bead-In-EMulsions (GEMs) via microfluidics. Within GEMs, polyadenylated mRNA transcripts were captured by oligo(dT)-linked gel beads, reverse-transcribed, and uniquely barcoded with cell- and molecule-specific identifiers (UMIs). Post-RT, GEMs were broken, and cDNA was amplified (12 cycles). Libraries were constructed by enzymatic fragmentation, end-repair, and adapter ligation, followed by sample indexing (10x Genomics Dual Index Kit) and PCR amplification (14 cycles). When passed QC, pooled libraries were sequenced on an Illumina Hiseq 4000, in the PE150 mode (OE Biotech Co., Ltd. Shanghai, China).

### The scRNA-seq data analysis on mouse heart specimens

Raw sequencing data were processed using Cell Ranger (10x Genomics, v8.0.1) on the mouse transcriptome reference of GRCm39 to generate gene-cell expression matrix files for nine specimens across three treatments (Sham, Model, and AH001). Expression matrix files for nine specimens were read and merged into a Seurat object using Seurat (v5.2.1) in R 4.4.3, with the filter out criteria of cells <3 and UMIs < 200. The merged Seurat object was processed by normalization, variable feature selection (top 2000), scaling, PCA dimensionality reduction, UMAP dimensionality reduction, and clustering (FindNeighbors and FindClusters). Cell population identity was assigned based on cell-type-specific markers to give 13 different cell types, using the SingleR (v 2.8.0) with the Mouse RNAseq Data reference dataset.^45^ Then, cardiomyocytes and fibroblasts were separated for downstream analysis. Differential gene expression (DGE) analysis was performed between different treatments in cardiomyocytes or fibroblasts. Ligand-receptor interactions between cardiomyocytes and fibroblasts were inferred using CellChat (v2.2.0) with the default mouse database.^46, 47^

### Data analysis on public datasets from human heart specimens

Gene-cell expression matrix files for the CITE-seq and H5 files for the snRNA-seq were downloaded from the GEO database, with the accession code of GSE218392.^19^ Expression matrix files were read and merged into a Seurat object using Seurat (v5.2.1) in R 4.4.3 with the filter out criteria of cells <3 and UMIs < 200. H5 files were converted into Seurat objects and merged into a Seurat object with the same filter out criteria. Cardiomyocytes and fibroblasts were separated according to the cell type annotation in the metadata file from the GEO database. DGE analysis was performed between Donors and patients in cardiomyocytes or fibroblasts. Gene sets for RHO GTPASE signaling were extracted from the REACTOME pathway database. The activation levels of RHO GTPASE signaling were evaluated in cardiomyocytes and fibroblasts using irGSEA (v2.1.5) ^48^ in R 4.4.3, with default parameters.

### Statistical analysis

For experimental data, intergroup comparisons were conducted using one-way ANOVA with LSD post hoc tests in GraphPad Prism 8.0 (GraphPad Software). Sequencing data analysis was performed with Seurat v5.2.1 in R 4.4.3, where statistical significance for intergroup comparisons on gene expressions was determined using adjusted p-values.

## SUPPLEMENTAL INFORMATION

**Figure S1 is related to Figure 1. Gene-level associations for RHOA and the baseline characteristics of heart failure patients and donors.**

(A and B) Common variant (A) and rare variant (B) gene-level association for RHOA comes from the Cardiovascular Disease Knowledge Portal (https://cvd.hugeamp.org/).

(C) The baseline characteristics of heart failure patients and donors giving interventricular septum (IVS) specimens. Etiologies from patients contained coronary artery disease (CAD) and atrial fibrillation (AF).

**Figure S2 is related to Figure 2. The discovery of a cryptic pocket and the impact of AH001 on the binding of RhoA to RhoGDIα.**

(A-C) The conformational landscape of RhoA ensembles from 600 ns MD simulations generated using the distance values between Tyr34 Cα atom in Switch I and Gly14 Cα atom in the P loop, and between Q63 Cα atom in Switch II and Gly14 Cα atom in the P loop. The crystal structures of RhoA^G14V^-GTPγS (PDB code: 1A2B in A) and RhoA^WT^- GDP (PDB code: 1FTN in B) were all projected onto the landscape (C).

(D) Comparison of the AH001-bound RhoA with apo RhoA (The first RhoA-apo structure in the asymmetric unit). The SW-I (29-42 aa), SW-II (58-73 aa) are colored cyan for RhoA-AH001, orange for RhoA-apo.

(E and F) The determination of GTP/GDP ratios by normalized F488 nm/F405 nm in (E, n = 3) with the representative images in (F). Scale bar, 20 μm.

(G) Comparison of RhoA-apo to RhoA-AH001-RhoGDIα complex. The structure of RhoA-AH001-RhoGDIα is derived from homology modeling based on the RhoA-RhoGDIα complex (PDB:1CC0).

(H) The schema of deletion of the regulator arm (5-55 aa) of RhoGDIα.

(I) PLA assays to detect the interaction between RhoA, RhoGDIα, and the truncated RhoGDIα in HEK 293F cells treated with AH001 at 20 μM, with the quantification of PLA signals. Scale bar, 5 μm.

Data are mean ± SEM. Statistical analysis was performed using one-way analysis of variance (ANOVA) among multiple groups, which was followed by an LSD post hoc test between two groups.

**Figure S4 is related to Figure 3. Impacts of AH001 on Ang II induced mouse models and determinations of cardiac injury indicators in LAD mouse models and rat models.**

(A) The schema of Ang II mouse models (n = 8) construction, followed by treatments of LCZ696 (66.67 mg/kg) and AH001 (10 mg/kg).

(B) EF and FS in Ang II mouse models by the Echocardiographic assessment.

(C-F) Representative images of HE, Masson’s trichrome staining, and WGA staining of a cross-sectional heart specimen, shown in the left ventricle region (C and E). Scale bar, 1 mm. Cardiac fibrosis was assessed by the fibrosis area and the cross-sectional area of cardiomyocytes (D and F).

(G) SBP and DBP determinations in Ang II mouse models.

(H) LDH, CK, and Ang II in serum from LAD mouse models were determined by ELISA assays.

LDH, CK, and BNP in serum from LAD rat models determined by ELISA assays. Data are mean ± SEM. Statistical analysis was performed using one-way analysis of variance (ANOVA) among multiple groups, which was followed by an LSD post hoc test between two groups.

**Figure S5 is related to Figure 3. Impacts of AH001 on Doxorubicin-induced mouse models.**

(A) The schema of Doxorubicin (Dox)-induced mouse models construction, followed by treatments of LCZ696 (60 mg/kg) and AH001 at doses of 1, 10, and 50 mg/kg.

(B) The survival of Dox-induced mice (n = 10) followed by treatments described in (A). (C and D) Body weight (C) and heart index (heart weight/body weight, D) in Dox-induced mouse models (n = 9).

(E) EF and FS in Dox-induced mouse models (n = 9) by the Echocardiographic assessment.

(F-H) LDH (F), HBDH (G), and CK (H) in serum from Dox-induced mouse models (n = 5) determined by ELISA assays.

(I-K) Representative images of HE staining (I) on heart specimens with longitudinal section (J) and transverse section (K). Cardiac vacuoles were indicated by arrows. Scale bar, 1mm (I), 20 μm (J and K).

(L) Representative images of Sirius red staining on heart specimens, with quantification of Sirius red^+^ area. Scale bar, 50 μm.

Data are mean ± SEM. Statistical analysis was performed using one-way analysis of variance (ANOVA) among multiple groups, which was followed by an LSD post hoc test between two groups.

**Figure S6 is related to Figure 5. The impact of secreted proteins from cardiomyocytes on fibroblasts.**

(A) Representative images of migration of fibroblasts in wound healing assays, with the treatments of AH001 at doses of 10, 20, and 40 μM. Scale bar, 200 μm.

(B) Representative images of migration of fibroblasts in Transwell assays, with the treatments of AH001 at doses of 10, 20, and 40 μM. Scale bar, 200 μm.

(C) Expression levels of proteins involved in fibrosis determined by WB, including FN1, COL3, SRF, VIM, and p-CFN.

(D) Representative images of immunofluorescence staining against ACTN2 in cardiomyocytes treated by AH001 at doses of 10, 20, and 40 μM, with quantification of the fluorescent area of ACTN2. Scale bar, 20 μm.

(E) Expression levels of proteins involved in fibrosis determined by WB, including FN1, COL3, SRF, and VIM.

(F) Representative images of immunofluorescence staining against ACTN2 in cardiomyocytes followed by the knockdown of RhoGDIα and the treatment of AH001 at 20 μM, with quantification of the fluorescent area of ACTN2. Scale bar, 20 μm.

(G) The schema of the AC-induced cardiomyocyte in vitro culture model construction, followed by treatments of AH001 at doses of 10, 20, and 40 μM.

(H) Determinations of secreted pro-fibrotic factors in the culture supernatant determined by ELISA assays.

(I) The impact of the conditioned medium from cardiomyocyte culture on the migration of fibroblasts (n = 3) as assessed by the Transwell assay. Scale bar, 200 μm.

Data are mean ± SEM. Statistical analysis was performed using one-way analysis of variance (ANOVA) among multiple groups, which was followed by an LSD post hoc test between two groups.

