## Supplemental Figures for "Inhibiting RhoA Activation via GDP-state Stabilization to Relieve Heart Failure"

A

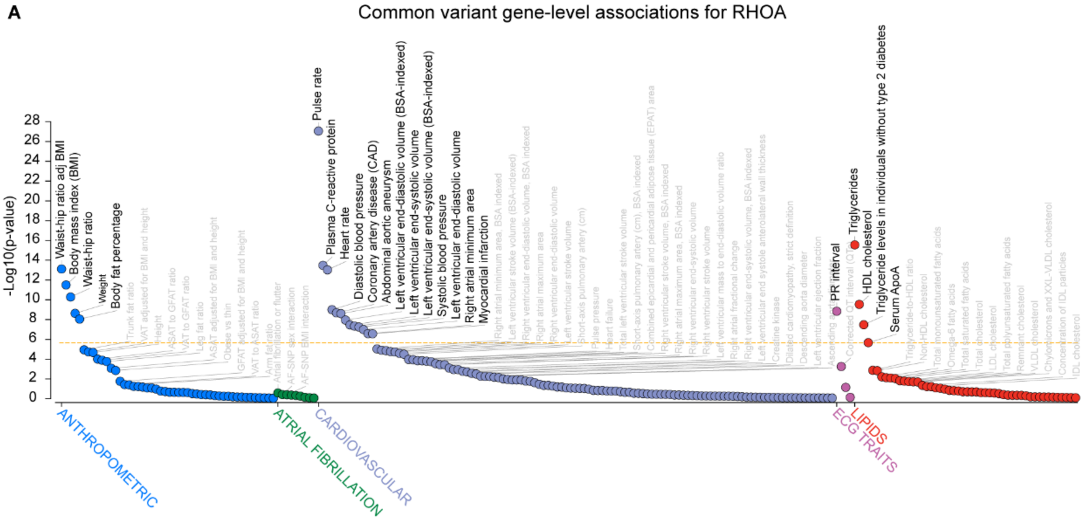


B

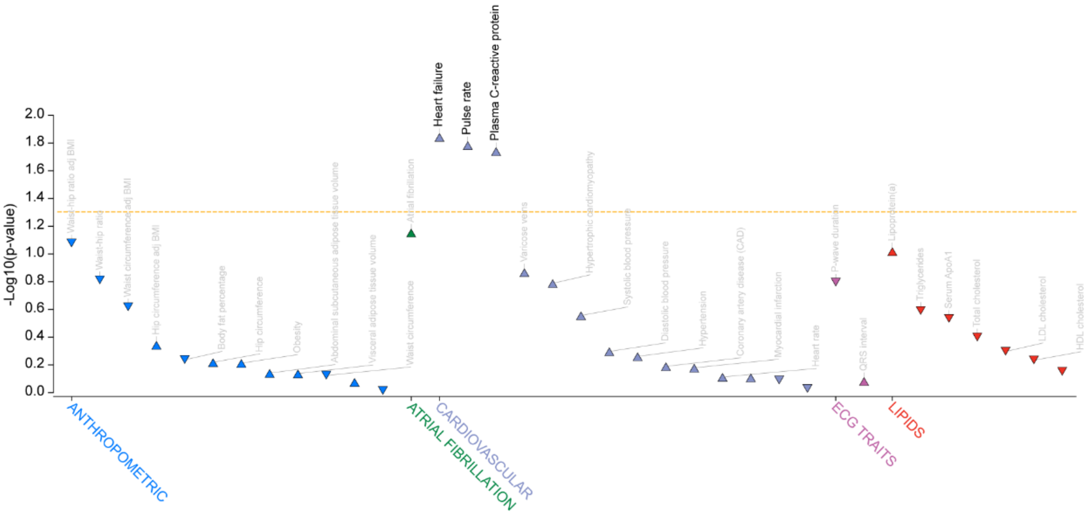


C

| ID | Age | Sex | Etiology | Specimen | NT-proBNP |
|---|---|---|---|---|---|
| Patient1 | 63 | Male | CAD | IVS | 242 |
| Patient2 | 58 | Male | AF | IVS | 185 |
| Patient3 | 69 | Female | CAD | IVS | 163 |

| ID | Age | Sex | Specimen |
|---|---|---|---|
| Donor1 | 38 | Male | IVS |
| Donor2 | 61 | Male | IVS |
| Donor3 | 52 | Female | IVS |

**Figure S1 is related to Figure 1. Gene-level associations for RHOA and the baseline characteristics of heart failure patients and donors.**

(A and B) Common variant (A) and rare variant (B) gene-level association for RHOA comes from the Cardiovascular Disease Knowledge Portal (https://cvd.hugeamp.org/).

(C) The baseline characteristics of heart failure patients and donors giving interventricular septum (IVS) specimens. Etiologies from patients contained coronary artery disease (CAD) and atrial fibrillation (AF).

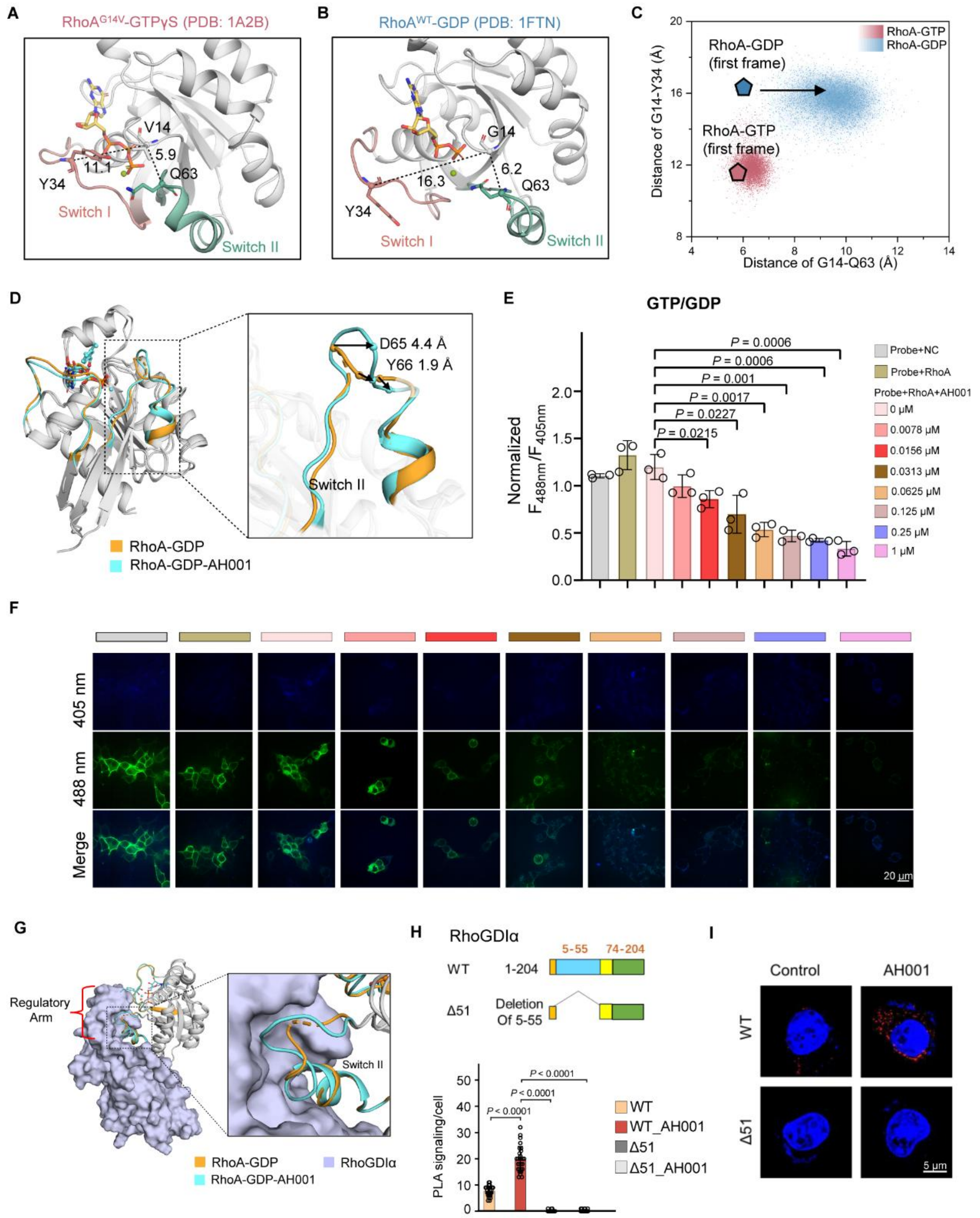


**Figure S2 is related to Figure 2. The discovery of a cryptic pocket and the impact of AH001 on the binding of RhoA to RhoGDIα.**

(A-C) The conformational landscape of RhoA ensembles from 600 ns MD simulations generated using the distance values between Tyr34 Cα atom in Switch I and Gly14 Cα atom in the P loop, and between Q63 Cα atom in Switch II and Gly14 Cα atom in the P loop. The crystal structures of RhoA$^{G14V}$-GTPγS (PDB code: 1A2B in A) and RhoA$^{WT}$-GDP (PDB code: 1FTN in B) were all projected onto the landscape (C).

(D) Comparison of the AH001-bound RhoA with apo RhoA (The first RhoA-apo structure in the asymmetric unit). The SW-I (29-42 aa), SW-II (58-73 aa) are colored cyan for RhoA-AH001, orange for RhoA-apo.

(E and F) The determination of GTP/GDP ratios by normalized F488 nm/F405 nm in (E, n = 3) with the representative images in (F). Scale bar, 20 μm.
(G) Comparison of RhoA-apo to RhoA-AH001-RhoGDIα complex. The structure of RhoA-AH001-RhoGDIα is derived from homology modeling based on the RhoA-RhoGDIα complex (PDB:1CC0).
(H) The schema of deletion of the regulator arm (5-55 aa) of RhoGDIα.
(I) PLA assays to detect the interaction between RhoA, RhoGDIα, and the truncated RhoGDIα in HEK 293F cells treated with AH001 at 20 μM, with the quantification of PLA signals. Scale bar, 5 μm.
Data are mean ± SEM. Statistical analysis was performed using one-way analysis of variance (ANOVA) among multiple groups, which was followed by an LSD post hoc test between two groups.

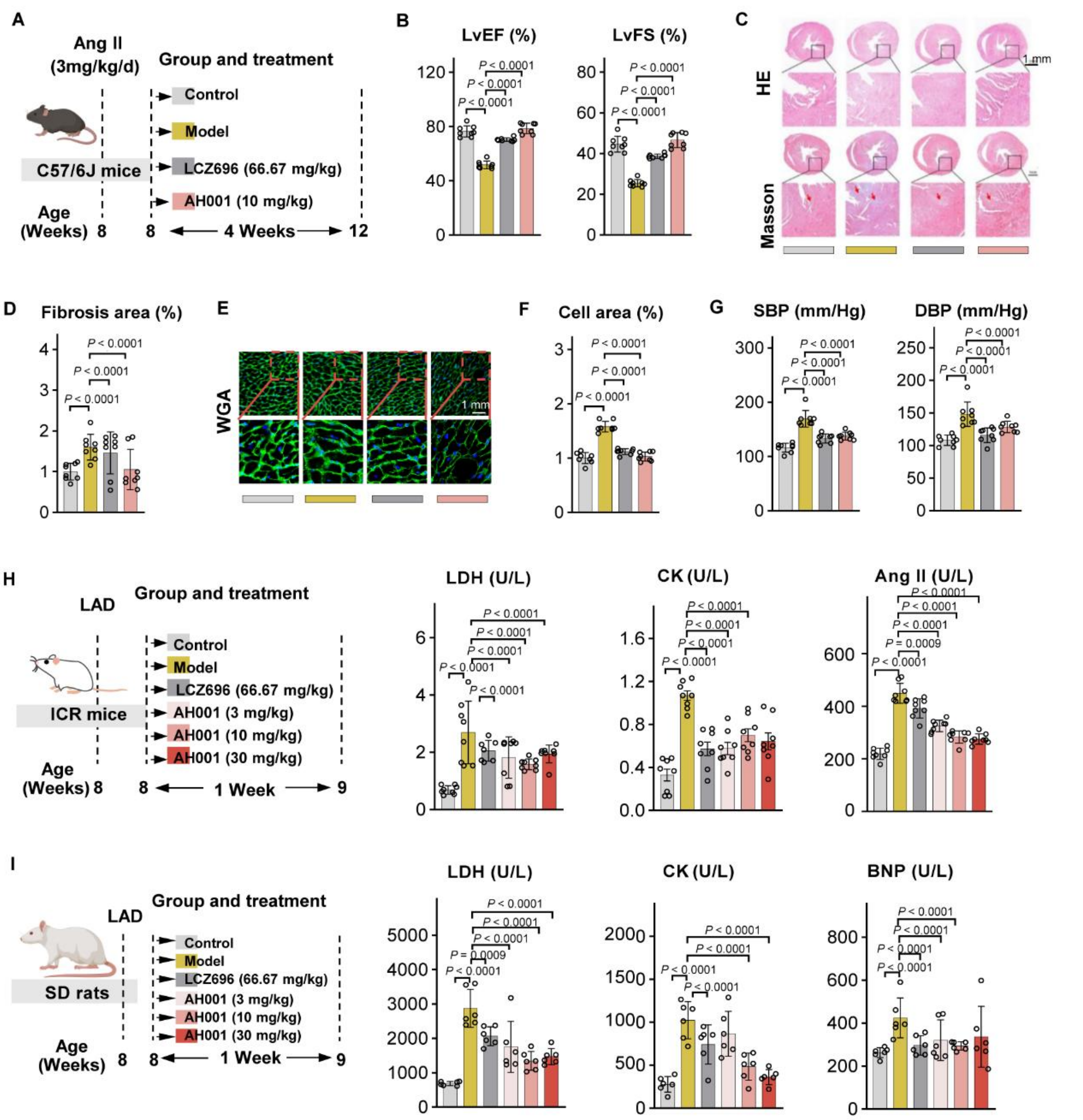


**Figure S4 is related to Figure 3. Impacts of AH001 on Ang II induced mouse models and determinations of cardiac injury indicators in LAD mouse models and rat models.**

(A) The schema of Ang II mouse models (n = 8) construction, followed by treatments of LCZ696 (66.67 mg/kg) and AH001 (10 mg/kg).

(B) EF and FS in Ang II mouse models by the Echocardiographic assessment.

(C-F) Representative images of HE, Masson's trichrome staining, and WGA staining of a cross-sectional heart specimen, shown in the left ventricle region (C and E). Scale bar, 1 mm. Cardiac fibrosis was assessed by the fibrosis area and the cross-sectional area of cardiomyocytes (D and F).

(G) SBP and DBP determinations in Ang II mouse models.

(H) LDH, CK, and Ang II in serum from LAD mouse models were determined by ELISA assays.

(I) LDH, CK, and BNP in serum from LAD rat models determined by ELISA assays.

Data are mean ± SEM. Statistical analysis was performed using one-way analysis of variance (ANOVA) among multiple groups, which was followed by an LSD post hoc test between two groups.

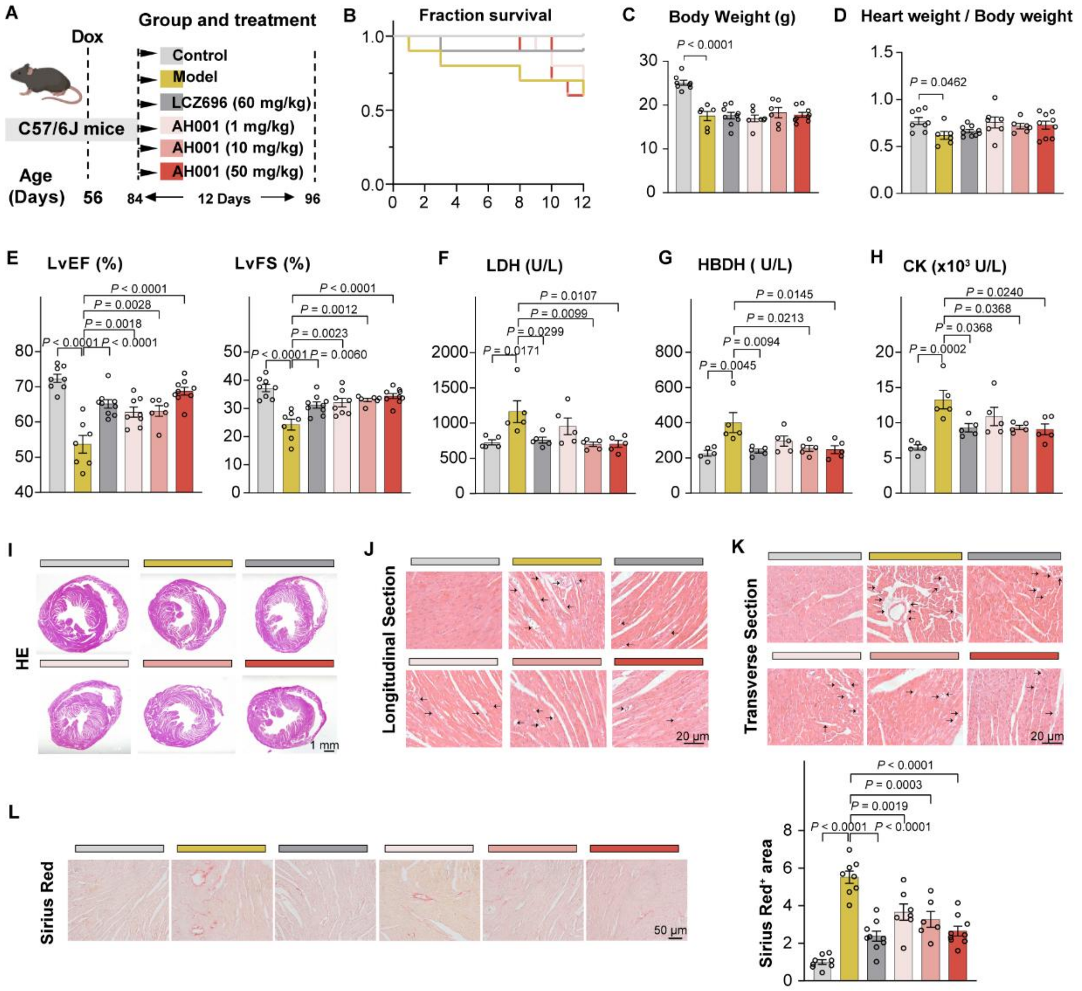


**Figure S5 is related to Figure 3. Impacts of AH001 on Doxorubicin-induced mouse models.**

(A) The schema of Doxorubicin (Dox)-induced mouse models construction, followed by treatments of LCZ696 (60 mg/kg) and AH001 at doses of 1, 10, and 50 mg/kg.

(B) The survival of Dox-induced mice (n = 10) followed by treatments described in (A).

(C and D) Body weight (C) and heart index (heart weight/body weight, D) in Dox-induced mouse models (n = 9).

(E) EF and FS in Dox-induced mouse models (n = 9) by the Echocardiographic assessment.

(F-H) LDH (F), HBDH (G), and CK (H) in serum from Dox-induced mouse models (n = 5) determined by ELISA assays.

(I-K) Representative images of HE staining (I) on heart specimens with longitudinal section (J) and transverse section (K). Cardiac vacuoles were indicated by arrows. Scale bar, 1mm (I), 20 μm (J and K).

(L) Representative images of Sirius red staining on heart specimens, with quantification of Sirius red$^{+}$ area. Scale bar, 50 μm.

Data are mean ± SEM. Statistical analysis was performed using one-way analysis of variance (ANOVA) among multiple groups, which was followed by an LSD post hoc test between two groups.

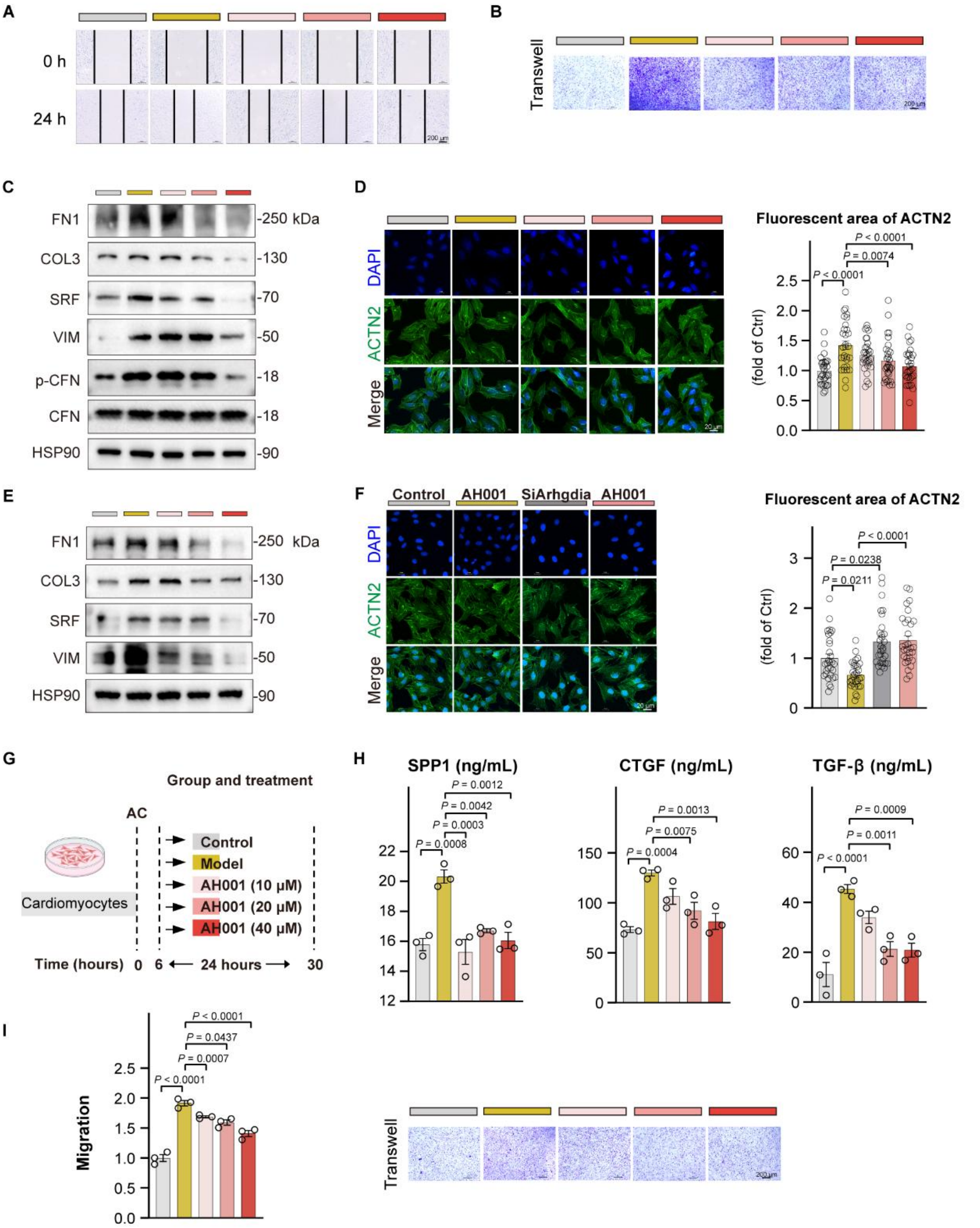


**Figure S6 is related to Figure 5. The impact of secreted proteins from cardiomyocytes on fibroblasts.**

(A) Representative images of migration of fibroblasts in wound healing assays, with the treatments of AH001 at doses of 10, 20, and 40 μM. Scale bar, 200 μm.

(B) Representative images of migration of fibroblasts in Transwell assays, with the treatments of AH001 at doses of 10, 20, and 40 μM. Scale bar, 200 μm.

(C) Expression levels of proteins involved in fibrosis determined by WB, including FN1, COL3, SRF, VIM, and p-CFN.

(D) Representative images of immunofluorescence staining against ACTN2 in cardiomyocytes treated by AH001 at doses of 10, 20, and 40 μM, with quantification of the fluorescent area of

ACTN2. Scale bar, 20 μm.
(E) Expression levels of proteins involved in fibrosis determined by WB, including FN1, COL3, SRF, and VIM.
(F) Representative images of immunofluorescence staining against ACTN2 in cardiomyocytes followed by the knockdown of RhoGDIα and the treatment of AH001 at 20 μM, with quantification of the fluorescent area of ACTN2. Scale bar, 20 μm.
(G) The schema of the AC-induced cardiomyocyte in vitro culture model construction, followed by treatments of AH001 at doses of 10, 20, and 40 μM.
(H) Determinations of secreted pro-fibrotic factors in the culture supernatant determined by ELISA assays.
(I) The impact of the conditioned medium from cardiomyocyte culture on the migration of fibroblasts (n = 3) as assessed by the Transwell assay. Scale bar, 200 μm.
Data are mean ± SEM. Statistical analysis was performed using one-way analysis of variance (ANOVA) among multiple groups, which was followed by an LSD post hoc test between two groups.
